# Haplotype-aware variant calling enables high accuracy in nanopore long-reads using deep neural networks

**DOI:** 10.1101/2021.03.04.433952

**Authors:** Kishwar Shafin, Trevor Pesout, Pi-Chuan Chang, Maria Nattestad, Alexey Kolesnikov, Sidharth Goel, Gunjan Baid, Jordan M. Eizenga, Karen H. Miga, Paolo Carnevali, Miten Jain, Andrew Carroll, Benedict Paten

## Abstract

Long-read sequencing has the potential to transform variant detection by reaching currently difficult-to-map regions and routinely linking together adjacent variations to enable read based phasing. Third-generation nanopore sequence data has demonstrated a long read length, but current interpretation methods for its novel pore-based signal have unique error profiles, making accurate analysis challenging. Here, we introduce a haplotype-aware variant calling pipeline PEPPER-Margin-DeepVariant that produces state-of-the-art variant calling results with nanopore data. We show that our nanopore-based method outperforms the short-read-based single nucleotide variant identification method at the whole genome-scale and produces high-quality single nucleotide variants in segmental duplications and low-mappability regions where short-read based genotyping fails. We show that our pipeline can provide highly-contiguous phase blocks across the genome with nanopore reads, contiguously spanning between 85% to 92% of annotated genes across six samples. We also extend PEPPER-Margin-DeepVariant to PacBio HiFi data, providing an efficient solution with superior performance than the current WhatsHap-DeepVariant standard. Finally, we demonstrate *de novo* assembly polishing methods that use nanopore and PacBio HiFi reads to produce diploid assemblies with high accuracy (Q35+ nanopore-polished and Q40+ PacBio-HiFi-polished).

## Introduction

Most existing reference-based small variant genotyping methods are tuned to work with short-reads [1, 2]. Short-reads have high base-level accuracy but frequently fail to align unambiguously in repetitive regions [3]. Short-reads are also generally unable to provide substantial read-based phasing information, and therefore require using haplotype panels for phasing [4] that provide limited phasing information for rarer variants.

Third-generation sequencing technologies, like linked-reads [5, 6, 7] and long-reads [8, 9], produce sequences that can map more confidently in the repetitive regions of the genome [10], overcoming the fundamental limitations of short-reads. Long-reads can generate highly contiguous *de novo* assemblies [11, 12, 13, 14, 15, 16, 17], and they are increasingly being used by reference-based analysis methods [18, 19, 20, 21, 22, 23, 24, 25]. The Genome-In-A-Bottle Consortium (GIAB) [26] used the additional power of long-reads and linked-reads to expand the small variant benchmarking set to cover more of the genome [27]. This was essential to the PrecisionFDA challenge V2, which quantified the limitations of short read-based methods to accurately identify small variants in repetitive regions[28].

Oxford Nanopore Technologies (ONT) is a commercial nanopore-based high-throughput [13] long-read sequencing platform that can generate 100kb+ long reads [13, 29]. Nanopore long-reads can confidently map to repetitive regions of the genome [10] including centromeric satellites, acrocentric short arms, and segmental duplications [11, 30, 31, 32]. Nanopore sequencing platform promises same-day sequencing and analysis [33], but the base-level error characteristics of the nanopore-reads, being both generally higher and systematic, make small variant identification challenging [34].

Pacific Biosciences (PacBio) provides a single-molecule real-time (SMRT) sequencing platform that employs circular consensus sequencing (CCS) to generate highly-accurate (99.8%) high-fidelity (PacBio HiFi) reads that are between 15kb-20kb long [21]. The overall accuracy of PacBio-HiFi-based variant identification compete with short-read based methods [21]. These highly accurate long-reads enabled the small variant benchmarking of major histocompatibility complex (MHC) region [35] and difficult-to-map regions [27].

In our previous work, we introduced DeepVariant, a universal small variant calling method based on a deep convolutional neural network (CNN) [36]. We showed that by retraining the neural network of DeepVariant, we can generate highly accurate variant calls for various sequencing platforms [36]. DeepVariant is a large neural network which can learn complex rules, to limit the computational time, DeepVariant only uses the neural network on candidate sites identified with simple heuristics. However, the higher error-rate of nanopore-reads [13, 34] causes too many candidate variants to be picked up by the heuristic-based candidate finder of DeepVariant, limiting the extension of DeepVariant to nanopore sequencing platform.

Previously, we trained DeepVariant on PacBio HiFi long-read data and showed highly competitive performance against short-read based methods for small variant identification [21]. We showed that using phasing information during genotyping with PacBio HiFi reads improved the overall genotyping accuracy [21]. However, the run-time of the haplotype-aware mode of DeepVariant with PacBio HiFi reads remain a bottleneck for production-level scaling.

Sufficiently accurate nanopore long-read based accurate small variant identification would enable new research. It could allow same-day sequencing and variant calling by using highly multiplexed sequencing with the PromethION device. It could allow researchers to study genomic variants in the most difficult regions of the genome. Similarly, making PacBio HiFi haplotype-aware genotyping efficient would allow researchers to adopt to production scale haplotype-aware genotyping.

Here we present a haplotype-aware genotyping pipeline PEPPER-Margin-DeepVariant that produces state-of-the art small variant identification results with nanopore and PacBio HiFi long-reads. PEPPER-Margin-DeepVariant outperforms other existing nanopore-based variant callers like Medaka[18], Clair [19], and longshot [20]. For the first time we report that nanopore-based single nucleotide polymorphism (SNP) identification with PEPPER-Margin-DeepVariant outperforms short-read based SNP identification with DeepVariant at whole genome scale. For PacBio HiFi reads, we report PEPPER-Margin-DeepVariant is more accurate and 3× faster and 1.4× cheaper than the current haplotype-aware pipeline DeepVariant-WhatsHap-DeepVariant. We analyzed our pipeline in the context of GENCODE[37] genes and report phasing errors in less than 1.5% of genes and over 88% of all genes being contiguously phased across six samples. Finally, we extended PEPPER-Margin-DeepVariant to polish nanopore-based *de novo* assemblies with nanopore and PacBio HiFi reads in a diploid manner. We report Q35+ nanopore-based and Q40+ PacBio-HiFi-polished assemblies with lower switch error rate compared to the unpolished assemblies.

## Results

### Haplotype-aware variant calling

PEPPER-Margin-DeepVariant is a haplotype-aware pipeline for identifying small variants against a reference genome with long-reads. The pipeline employs several methods to generate highly-accurate variant calls (Figure 1a). Details of these methods are in the online methods section. An overview is presented here:

1. **PEPPER-SNP**: PEPPER-SNP finds single nucleotide polymorphisms (SNPs) from the read alignments to the reference using a recurrent neural network (RNN). The method works in three steps:

- *Image generation:* We take the reads aligned to a reference genome and generate base-level summary statistics in a matrix-like format for each location of the genome. We do not encode insertions observed in reads at this stage.
- *Inference:* We use a gated recurrent unit (GRU)-based RNN that takes the base-level statistics generated in the previous step and the provides likelihood of the two most likely bases present at each genomic location.
- *Find candidates:* We take all of the base-mismatches observed in the reads-to-reference alignment. We calculate a likelihood using the predictions from the inference step and if a base-mismatch has high likelihood of being a potential variant, we pick the mismatch to be a potential SNP. The likelihood of the bases at any location also helps to assign a genotype.
2. **Margin**: Margin is a phasing and haplotyping method that takes the SNPs reported by PEPPER-SNP and generates a haplotagged alignment file using a hidden Markov Model (HMM).

- *Read-allele alignment*: We first extract read substrings around allelic sites and generate alignment likelihoods between reads and alleles. These are used as emission probabilities in the phasing HMM.
- *Phasing Variants:* We construct an HMM describing genotypes and read bipartitions at each variant site which enforces consistent partitioning between sites. After running the forwardbackward algorithm, we marginalize over the posterior probability distribution at each site to calculate the most likely phased genotype (aka diplotype).
- *Haplotagging reads*: After determining haplotypes using the maximum probability haplotype decoding, we decide from which haplotype each read originated from by calculating the probability of the read arising from each of the two haplotypes and picking the haplotype with maximum likelihood. If a read spans no variants or has equal likelihood for each haplotype, it is assigned the a “not haplotagged” tag.
- *Chunking:* The genome is broken up into 120kb chunks with 20kb of overlap between chunks. Variant and read phasing occurs separately on each chunk, enabling a high degree of parallelism. Chunks are stitched together using the haplotype assignment of the reads shared between adjacent chunks.
3. **PEPPER-HP**: PEPPER-HP takes the haplotagged alignment file and finds potential SNP, insertion, and deletion (INDEL) candidate variants using a recurrent neural network (RNN). In this step PEPPER-HP ranks all variants arising from the read-to-reference alignment and picks variants with high-likelihood derived from the RNN output. Filtering candidates enables DeepVariant to efficiently genotype the candidates and produce a highly accurate variant set as it removes errors. PEPPER-HP is used only during Oxford Nanopore-based variant calling and has proved unnecessary while using PacBio HiFi reads.

- *Image generation:* We generate base-level summary statistics for each haplotype independently. Summary statistics for each haplotype use both reads that were haplotagged to that haplotype and which were not haplotagged. In this scheme, we encode insertions observed in the reads.
- *Inference:* We use a GRU-based RNN that takes the haplotype-specific summary statistics and predicts two bases at each location of the genome, one for each haplotype. This haplotype-aware inference scheme allows us to determine most likely alleles in a haplotype-specific manner.
- *Find candidates:* In the find candidates step we find all SNP and INDEL candidates arising from the read alignment to the reference. We use the haplotype-specific predictions from the inference step to generate the likelihood of each candidate variant belonging to haplotype-1 or haplotype-2. Using the likelihood values we propose candidates with high likelihood for genotyping with DeepVariant.
4. **DeepVariant**: DeepVariant identifies variants in a three step process:

- *Make examples:* Prior to this work [36], the *make examples* stage of DeepVariant used simple heuristics to identify possible variant positions for classification. The different error profile of Oxford Nanopore required the more sophisticated logic from PEPPER to generate a tractable number of candidates for classification. DeepVariant was modified to take the candidate variant set from PEPPER-HP and the haplotagged alignment from Margin, and to generate the tensor input set with read features as channels (base, base quality, mapping quality, strand, whether a read supports the variant, and the bases that mismatch the reference). Reads are sorted by their haplotype tag.
- *Call variants:* This stage applies a model trained specifically for Oxford Nanopore data with inputs provided by PEPPER-Margin. Apart from training on new data, and the sorting of reads by haplotype, other software components of this step are unchanged.
- *Postprocess variants:* Converts the output probability into a VCF call and resolves multi-allelic cases. No other changes were made from previously published descriptions. Combining PEPPER with DeepVariant in this way allows the faster and smaller neural network of PEPPER to scan much more of the genome, and for the larger but slower neural network of DeepVariant to apply more complicated models to a tractable number of candidates.
5. **Margin**: Margin takes the output of DeepVariant and the alignment file to generate a phased VCF file using the same Hidden Markov Model as described before. In this mode, it annotates the VCF with high-confidence phasesets using heuristics over the reads assigned to each variant’s haplotype. It creates a new phaseset if there is no linkage between adjacent sites, if there is an unlikely binomial p-value for the bipartition of reads at a site, or if there is high discordancy between read assignments over adjacent variants.

We trained PEPPER-Margin-DeepVariant on HG002 sample using Genome-In-A-Bottle (GIAB) v4.2.1 benchmarking set. We trained PEPPER and DeepVariant on `chr1-chr19` and tested on `chr20` and used `chr21-chr22` as holdout sets (See Online Methods).

### Nanopore variant calling performance

We compared the nanopore variant calling performance of PEPPER-Margin-DeepVariant against Medaka[18], Clair[19], and Longshot[20]. We called variants on two samples HG003 and HG004, with 90× coverage. We also compared the performance against Medaka and Clair for the HG003 sample at various coverages ranging from 20× to 90×. Finally, we benchmarked the variant calling performance of PEPPER-Margin-DeepVariant on six Genome-In-A-Bottle (GIAB) samples.

PEPPER-Margin-DeepVariant produces more accurate nanopore-based SNP calls (F1-scores of 0.9969 and 0.9977) for HG003 and HG004 respectively than Medaka (0.9926, 0.9933), Clair (0.9861, 0.9860), and Longshot (0.9775, 0.9776). We also observe higher INDEL performance with PEPPER-Margin-DeepVariant (F1-scores of 0.7257 and 0.7128 for HG003 and HG004) compared to Medaka (0.7089, 0.7128) and Clair (0.5352, 0.5260)(Figure 1b, Supplementary table 1).

**Figure 1:**
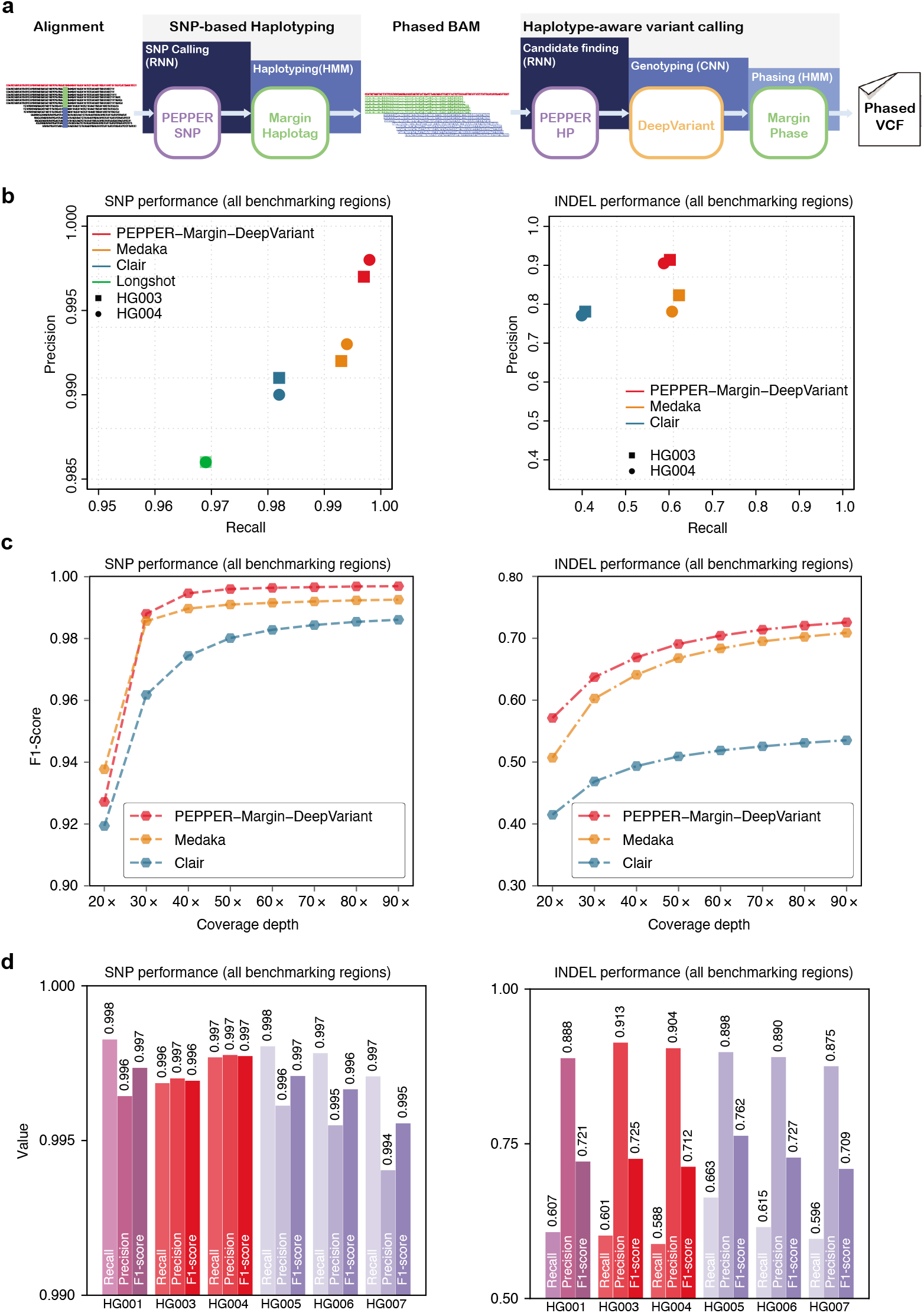
Nanopore variant calling results. **(a)** Illustration of haplotype-aware variant calling using PEPPER-Margin-DeepVariant. **(b)** Nanopore variant calling performance comparison between different nanopore-based variant callers. **(c)** Evaluating variant calling performance at different coverage of HG003. **(d)** Variant calling performance of PEPPER-Margin-DeepVariant on six GIAB samples.

To understand performance over realistic coverage ranges, we downsampled the HG003 nanopore sample at coverages varying between 20× and 90× and compared PEPPER Margin-DeepVariant against Medaka and Clair. The INDEL performance of PEPPER-Margin-DeepVariant achieves the highest F1-score at any coverage compared to other tools (Figure 1c, Supplementary table 2). At coverage above 30×, PEPPER-Margin-DeepVariant achieves a higher F1-score than Medaka and Clair (Supplementary table 3). Overall, we observe that PEPPER-Margin-DeepVariant can yield high-quality variant calls at above 40× coverage on Oxford Nanopore data.

We investigated the nanopore variant calling performance of PEPPER-Margin-DeepVariant on six GIAB samples (HG001, HG003-HG007), each sample with various coverage (Supplementary Table 4) and against GRCh37 and GRCh38 reference genomes (Supplementary Table 5). PEPPER-Margin-DeepVariant achieves SNP F1-score 0.995 or higher and INDEL F1-score of 0.709 or higher for each sample, demonstrating the ability to generalize the variant calling across samples and reference genomes (Figure 1d, Supplementary table 6).

We compared the run-time and cost of Oxford nanopore-based variant calling pipelines on 50× and 75× HG001 data (Table 1, Supplementary table 7). Clair (HG001-50×: 2.5h/$11.40, HG001-75×: 3.1h/$14.13) is the most run-time and cost efficient variant caller for Oxford Nanopore, however, previous analysis (Figure 1, Supplementary Table 1) show that Clair fails to generate high-quality variant calls. Longshot (HG001-50×: 51h/$49, HG001-75×: 74h/$139) is single-theraded and has higher memory requirement, which forces to use high-memory instances and increases the cost. Similarly, Medaka can not use all the available resources. Medaka can only use 2-CPUs but have a higher memory requirement which makes it unsuitable for CPU-based instances (HG001-50×: 95h/$90, HG001-75×: 117h/$175). We observed run-time improvement on a GPU platform with Medaka (HG001-50×: 40h/$97, HG001-75×: 46h/$225), but the memory requirement increases the cost. PEPPER-Margin-DeepVariant is designed for CPU and GPU platforms. On a CPU-platform, PEPPER-Margin-DeepVariant (HG001-50×: 13h/$60, HG001-75×: 15h/$68) is 8× faster than Medaka and 4× faster than Longshot while providing the best variant calling performance. On GPU-platforms we see further run-time improvement with PEPPER-Margin-DeepVariant (HG001-50×: 7h/$70, HG001-75×: 9h/$94). Overall, PEPPER-Margin-DeepVariant provides a scalable solution to haplotype-aware variant calling with nanopore-based long reads as it is designed to efficiently use all available resources.

**Table 1:**
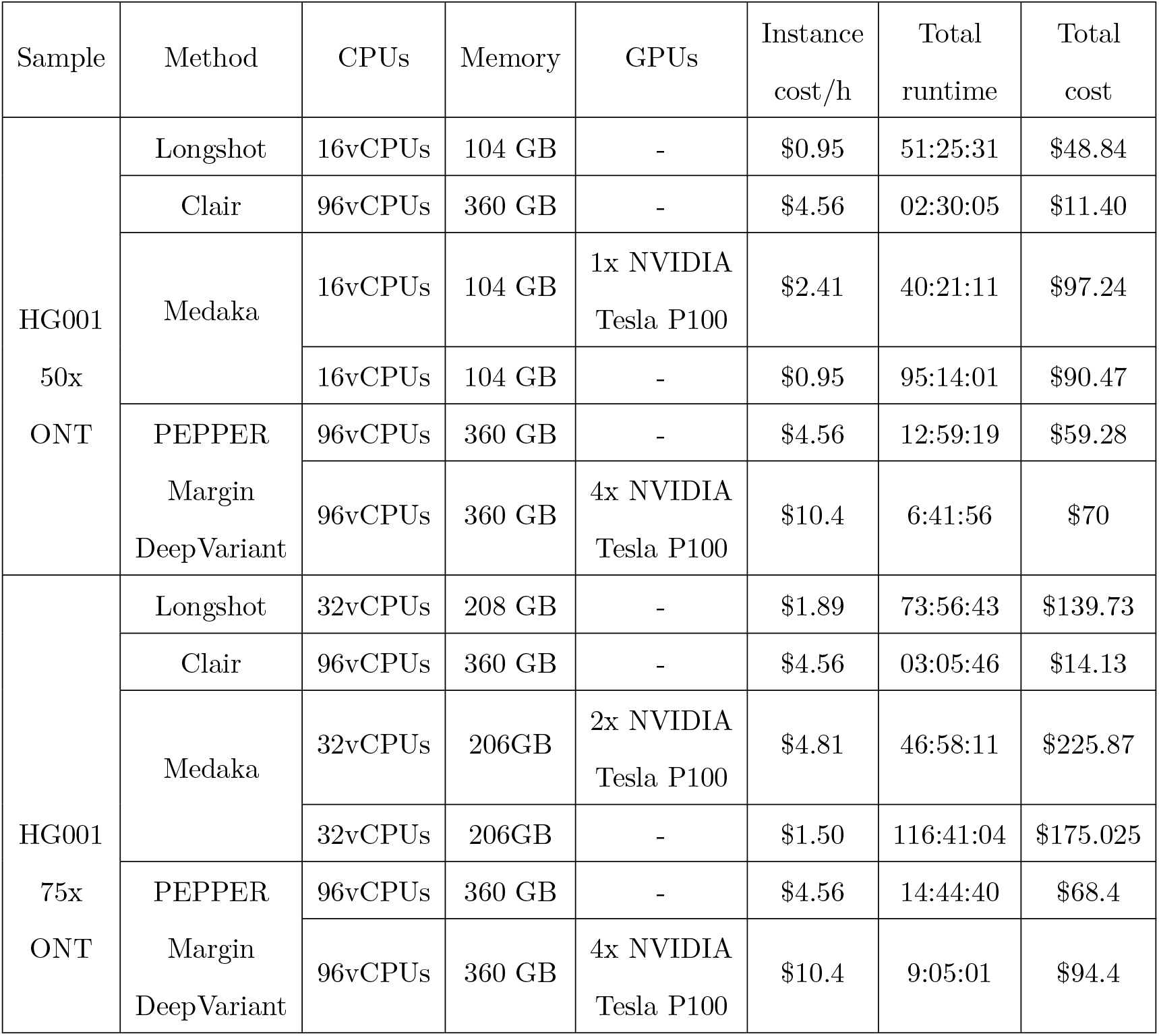
Run-time and cost analysis of Oxford nanopore-based variant calling pipelines on 50x and 75x HG001 data. We used various `n1-series` instance types available on Google Cloud Platform (GCP). The we calculated the cost using the GCP cost calculator (https://cloud.google.com/products/calculator). Logs of all the runs are publicly available (See supplementary Notes).

### PacBio HiFi variant calling performance

Similar to the nanopore-based haplotype-aware pipeline the PacBio HiFi based PEPPER-Margin-DeepVariant pipeline produces highly accurate variant calls. In the PacBio HiFi pipeline, we do not use PEPPER-HP to find candidate variants; the highly accurate (99.8%) PacBio HiFi reads are suitable for the heuristic-based candidate generation approach of DeepVariant [21, 36].

In table 2, we show the PacBio HiFi PEPPER-Margin-DeepVariant variant calling performance on the 35x HG003 and HG004 from precisionFDA [28] against DeepVariant-WhatsHap-DeepVariant and DeepVariant-Margin-DeepVariant where DeepVariant-WhatsHap-DeepVariant is currently the state-of-the-art method [28, 21]. In this comparison we see that DeepVariant-Margin-DeepVariant produces best performance (HG003 SNP-F1: 0.9991 INDEL-F1: 0.9945, HG004 SNP-F1: 0.9992, INDEL-F1: 0.9942) compared to DeepVariant-WhatsHap-DeepVariant (HG003 SNP-F1:0.9990 INDEL-F1:0.9942, HG004 SNP-F1: 0.9992, INDEL-F1: 0.9940) and PEPPER-Margin-DeepVariant (HG003 SNP-F1:0.9990, INDEL-F1:0.9944, HG004 SNP-F1: 0.9992, INDEL-F1: 0.9941). Notably, the PEPPER-Margin-DeepVariant pipeline outperforms DeepVariant-WhatsHap-DeepVariant and is 3× faster and 1.4× cheaper, which establishes a faster and more accurate solution to haplotype-aware variant calling with PacBio HiFi data (Supplementary table 8, Supplementary table 9).

**Table 2:**
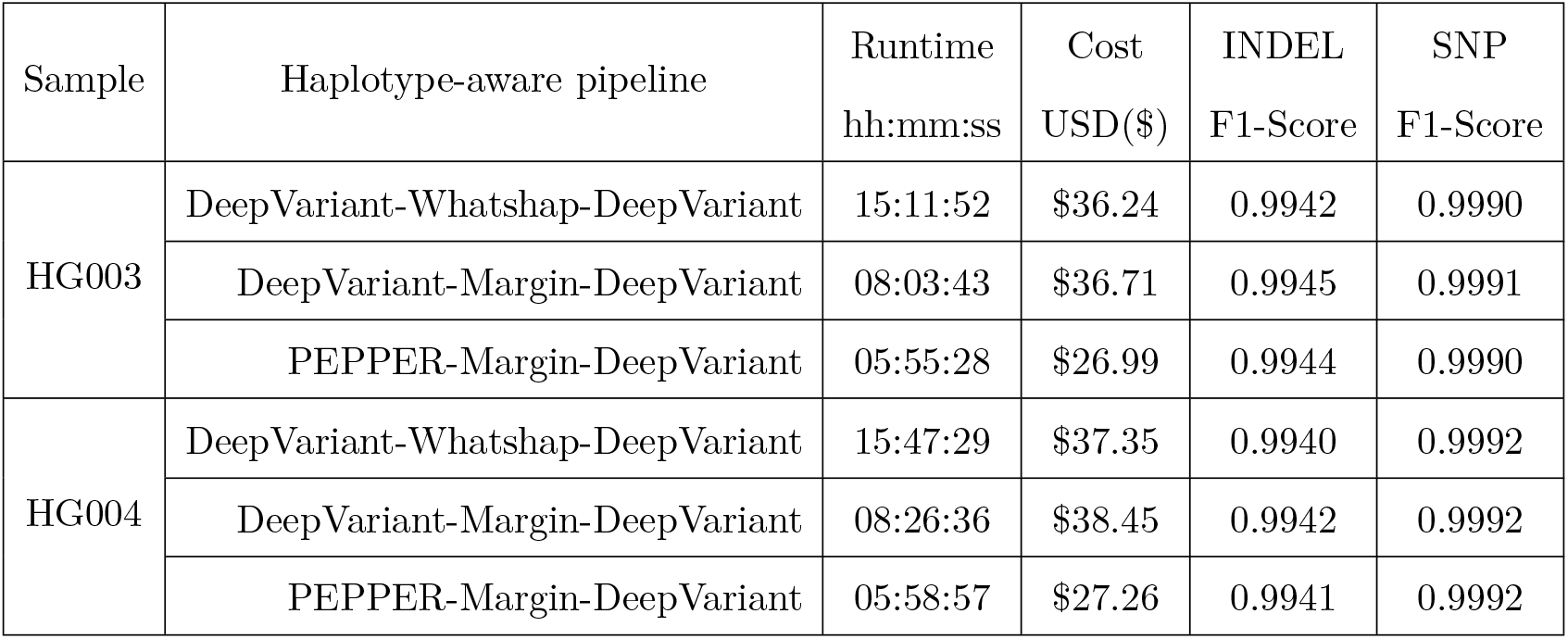
PacBio HiFi variant calling performance and runtime comparison between three haplotype-aware pipelines on 35 × coverage HG003 and HG004 samples. For PEPPER, Margin and DeepVariant we used $4.56/`h n1-standard-96` and for WhatsHap we used $0.09/`h n1-standard-2` instance types on google cloud platform. The F1-scores are derived by comparing the variant calls against GIAB v4.2.1 benchmark variants for HG003 and HG004.

| Sample | Haplotype-aware pipeline | Runtime | Cost | INDEL | SNP |
| --- | --- | --- | --- | --- | --- |
| | | hh:mm:ss | USD(\$) | F1-Score | F1-Score |
| HG003 | DeepVariant-WhatsHap-DeepVariant | 15:11:52 | \$36.24 | 0.9942 | 0.9990 |
| | DeepVariant-Margin-DeepVariant | 08:03:43 | \$36.71 | 0.9945 | 0.9991 |
| | PEPPER-Margin-DeepVariant | 05:55:28 | \$26.99 | 0.9944 | 0.9990 |
| HG004 | DeepVariant-WhatsHap-DeepVariant | 15:47:29 | \$37.35 | 0.9940 | 0.9992 |
| | DeepVariant-Margin-DeepVariant | 08:26:36 | \$38.45 | 0.9942 | 0.9992 |
| | PEPPER-Margin-DeepVariant | 05:58:57 | \$27.26 | 0.9941 | 0.9992 |

### Nanopore, Illumina and PacBio HiFi variant calling performance comparison

We compared the variant calling performance of Oxford Nanopore and PacBio HiFi long-read based PEPPER-Margin-DeepVariant against Illumina short-read based DeepVariant method. We used 35x Illumina NovaSeq, 35x PacBio HiFi, and 90x Oxford Nanopore reads basecalled with Guppy v4.2.2 for HG003 and HG004 samples available from PrecisionFDA [28]. We used GIAB v4.2.1 benchmarking data for HG003 and HG004, which is notable for including difficult-to-map regions. Finally we used GIAB v2.0 stratificiations to compare variant calling performance in difficult-to-map regions and low-complexity regions of the genome.

The SNP F1-score of PacBio HiFi (HG003 SNP-F1: 0.9990, HG004 SNP-F1: 0.9992) is higher than Oxford Nanopore (HG003 SNP-F1: 0.9969, HG004 SNP-F1: 0.9977) and Illumina (HG003 SNP-F1: 0.9963, HG004 SNP-F1: 0.9962) in all benchmarking regions. Notably, both long-read sequencing platforms outperform the short-read based method in accurate SNP identification performance. The INDEL F1-score of Oxford Nanopore (HG003 INDEL-F1: 0.7257, HG004 INDEL-F1: 0.7128) is well below the performance with PacBio HiFi (HG003 INDEL-F1: 0.9945, HG004 INDEL-F1: 0.9941) and Illumina (HG003 INDEL-F1: 0.9959, HG004 INDEL-F1: 0.9958) suggesting further improvement required for nanopore-based methods. Overall, we find that haplotype-aware long-read-based variant calling produces high-quality SNP variant calls comparable to those produced by short-read-based variant identification methods (Figure 2a, Supplementary table 10). This is the first demonstration we are aware of in which SNP variant calls with Oxford Nanopore data achieved similar accuracy to Illumina SNP variant calls.

**Figure 2:**
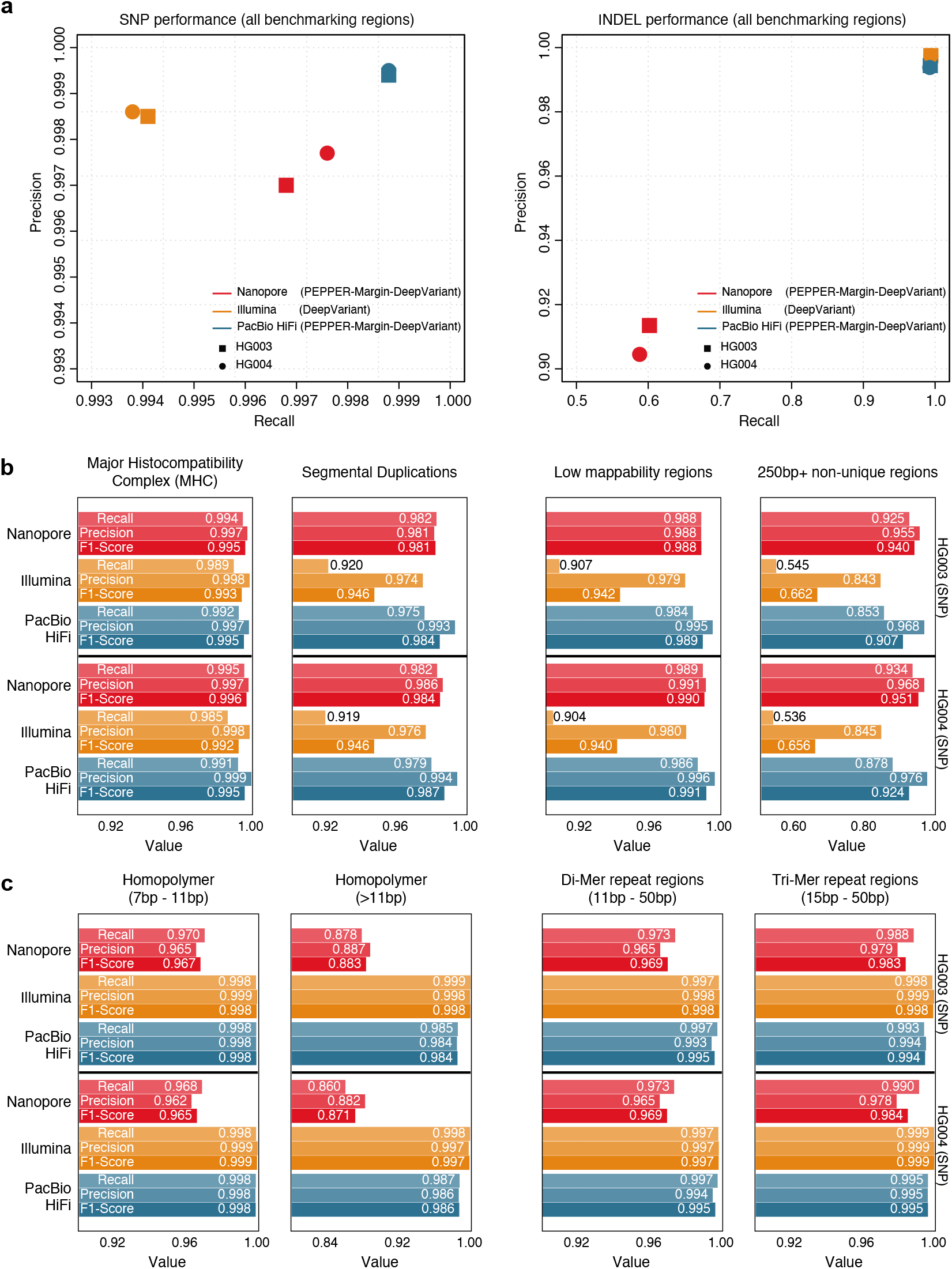
Comparison between Nanopore, Illumina and PacBio HiFi variant calling performance. **(a)** SNP and INDEL performance comparison of Nanopore, Illumina and PacBio HiFi in all benchmarking regions. **(b)** SNP performance comparison in difficult-to-map regions of the genome. **(c)** SNP performance comparison in low-complexity regions of the genome.

In segmental duplications, 250bp+ non-unique regions, and low-mappability regions where short-reads have difficulty in mapping, we observe the average SNP F1-scores of Illumina (Seg. Dup. F1-score: 0.94, 250bp+ non-unique:0.66, Low-mappability: 0.94) drop sharply for both HG003 and HG004 samples. Long-read based PacBio HiFi (Seg. Dup. F1-score: 0.99, 250bp+ non-unique:0.90, Low-mappability: 0.99) and Oxford Nanopore (Seg. Dup. F1-score: 0.98, 250bp+ non-unique:0.94, Low-mappability: 0.98) produce more accurate SNP variants. In the major histocompatibility complex (MHC) region, we see Oxford Nanopore (HG003 SNP F1-score: 0.9958, HG004 SNP F1-score: 0.9966) achieve best performance followed by PacBio HiFi (HG003 SNP F1-score: 0.9951, HG004 SNP F1-score: 0.9955) and Illumina HG003 SNP F1-score: 0.9939, HG004 SNP F1-score: 0.9921). In general, the long-read-based haplotype-aware methods outperform short-reads in more repetitive regions of the genome (Figure 2b, Supplementary table 11).

In low-complexity regions like homopolymer, di-mer and tri-mer repeat regions of the genome, the average variant calling performance of Nanopore drops (7bp-11bp homopolymer SNP F1-score: 0.96, 11bp+ homopolymer SNP F1-Score: 0.88) for both HG003 and HG004 samples compared to Illumina (7bp-11bp homopolymer SNP F1-score: 0.998, 11bp+ homopolymer SNP F1-Score: 0.998) and PacBio HiFi (7bp-11bp homopolymer SNP F1-score: 0.998, 11bp+ homopolymer SNP F1-Score: 0.984). In 11bp-50bp di-mer and 15bp-50bp tri-mer repeat regions of the genome, we see the average performance of Oxford Nanopore (di-mer SNP F1-score: 0.969, tri-mer SNP F1-score: 0.984) is lower than PacBio HiFi (di-mer SNP F1-score: 0.995, tri-mer SNP F1-score: 0.995) and Illumina (di-mer SNP F1-score: 0.998, tri-mer SNP F1-score: 0.998). Overall, the Illumina short-read based variant calling method achieves higher accuracy in low-complexity regions of the genome (Figure 2c, Supplementary table 12).

### Phaseset and Haplotagging Accuracy

We compared phaseset accuracy for Margin and WhatsHap on HG001 against GIAB’s phased v3.3.2 variants with 25× nanopore, 50× nanopore, 75× nanopore, and 35× PacBio HiFi data. We generated genotyped variants with PEPPER-Margin-DeepVariant, and used both Margin and WhatsHap to phase the final variant set. The phasesets produced by both tools were analyzed using `whatshap stats` and `whatshap compare` against the trio-confirmed truth variants in high-confidence regions.

For all datasets, Margin had a lower switch error rate (0.00875, 0.00857, 0.00816, 0.00895) than WhatsHap (0.00923, 0.00909, 0.00906, 0.00930), but lower phaseset N50 (2.07, 4.21, 6.13, 0.24 Mb) than WhatsHap (2.37, 4.90, 8.27, 0.25 Mb) (Figure 3a, Supplementary Tables 13 and 14).

**Figure 3:**
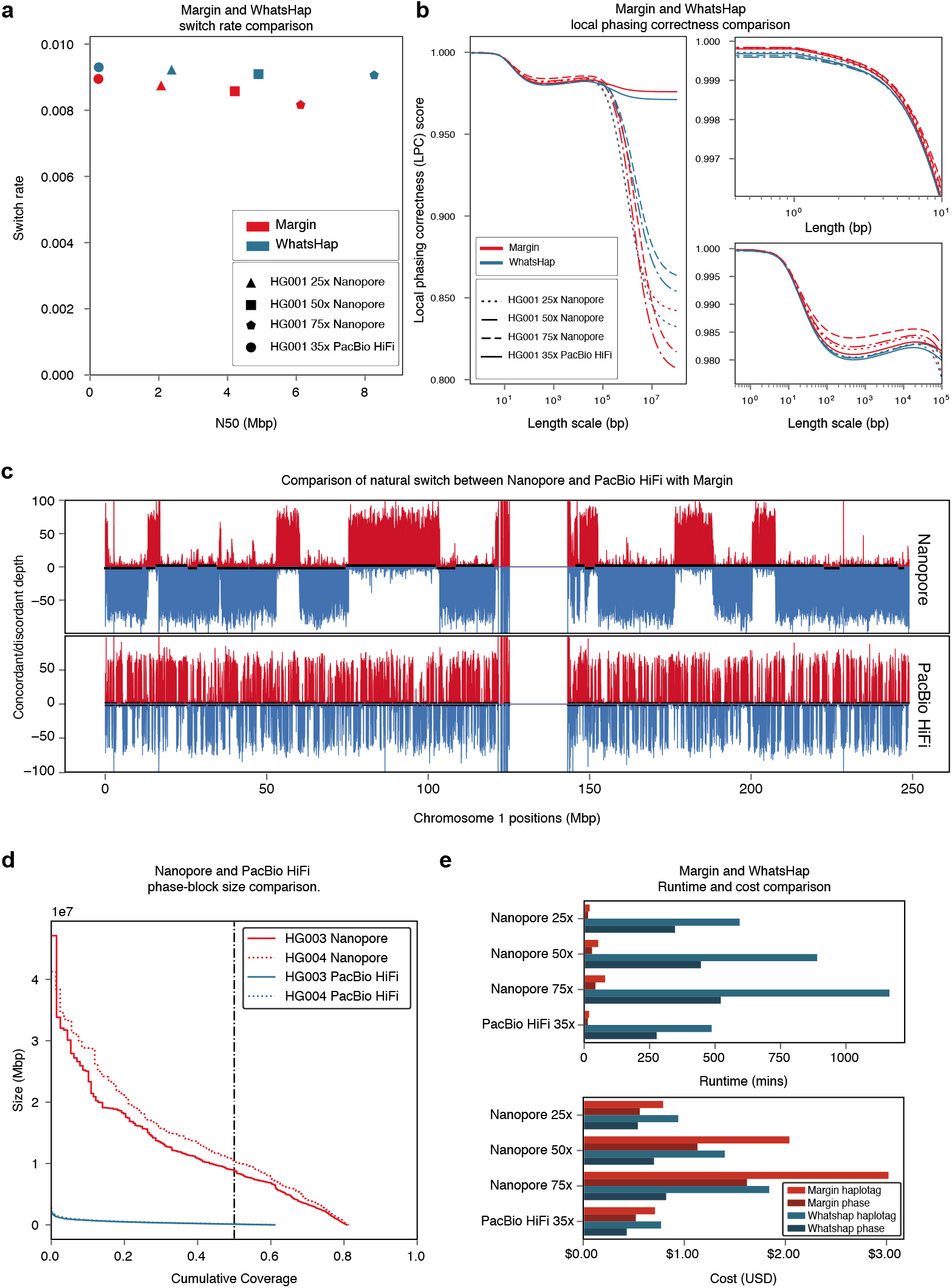
Margin and WhatsHap phasing results. **(a)** Phaseset switch rate to N50. **(b)** Novel metric “Local Phasing Correctness” analyzing phaseset accuracy across different length scales. For visibility, we plot the full data (left), short length scales (right top), and medium length scales (right bottom). **(c)** Novel visualization “Natural Switch Rate” describing haplotagging accuracy for reads. Phasing is consistent across the top/red and bottom/blue blocks, with switch errors occurring at the transitions between these. **(d)** Phaseset N50 for Nanopore and PacBio HiFi data on HG003 and HG004. **(e)** Cost and runtime comparison between Margin and Whatshap. Plots **(a)** and **(b)** are generated from phased HG001 GIAB v3.3.2 variants; plot **(c)** is generated from an admixture of reads from HG005 and HG02723.

We also compared phaseset accuracy for Margin and WhatsHap on the same data using a novel metric we call “Local Phasing Correctness” (LPC). In brief, the LPC is a value between 0 and 1 that summarizes whether every pair of heterozygous variants is correctly phased relative to each other. The contribution of each pair of variants is weighted based on the distance between them, with the weights varying according to a tunable parameter, the “length scale”. The length scale can be understood roughly as the scale of distances that influence the metric (see online methods). The LPC is a generalization of the standard metrics of switch error rate and Hamming rate, which have a close relationship with the LPC at length scales 0 and infinity respectively. We plot the LPC across various length scale values (Figure 3b). Margin produced more accurate phasing for all length scales for 25× nanopore and 35× CCS. Margin also produced more accurate phasing for 50× nanopore for length scales up to 128kb and for 75× nanopore for length scales up to 242kb, after which WhatsHap outperforms Margin. Both tools exhibit local maxima for length scales from 20-30 kilobases.

To analyze haplotagging accuracy, we artificially constructed an admixture sample by trio-binning reads from HG005 and HG02723 and combining an equal amount of maternal reads from each sample, resulting in a 55× nanopore alignment and a 35× PacBio HiFi alignment. We ran PEPPER-SNP, haplotagged each alignment with Margin using these variants, and compared the number of direct-matched reads ***R_d_*** (truth H1 to tagged H1 or truth H2 to tagged H2) and cross-matched reads ***R_c_*** (truth H1 to tagged H2 or truth H2 to tagged H1) of the output. In Figure 3c, for each 10kb bucket in chr1 we plot the number of reads that were direct-matched (top, red) and cross-matched (bottom, blue) for both data types, with phasesets plotted in black alternating between top and bottom. With this “Natural Switch” plot, it is possible to identify consistent phasing as regions where the majority of reads are either direct- or cross-matched, and switch errors in the haplotagging as regions where the majority of reads transition between the two. As the plot shows, nanopore reads allow us to haplotag consistently with phase sets in the range of tens of megabases, whereas PacBio HiFi reads cannot be used for long-range haplotagging. For each bucket we can calculate a local haplotagging accuracy using the ratio: max(***R_c_, R_d_***)/(***R_c_ + R_d_***). On average the haplotagging accuracy is 0.9557 for ONT data and 0.9764 for HiFi data using Margin (Supplementary Table 15). Full plots including local haplotagging accuracy visualization are shown in Supplementary Figures 2, 3, 4.

Lastly we compare the runtime and cost for the haplotag and phase actions on the four HG001 datasets using Margin and WhatsHap. When configured to use 64 threads, Margin at peak used 35GB of memory on a GCP instance `n1-highcpu-64` costing $2.27/hr. This results in a total cost (for haplotagging and phasing) of $1.35 (36m) for 25x ONT, $3.17 (84m) for 50x ONT, $4.64 (123m) for 75x ONT, $1.23 (33m) for 35x PacBio HiFi. Given WhatsHap’s concurrent use of two threads and three GB of memory we determined it could be run most cheaply on the GCP `n1-standard-2` instance type for $0.095/hr, resulting in a total cost of $1.48 (941m), $2.10 (1336m), $2.66 (1688m), and $1.20 (764m) respectively.

### Gene Analysis

We performed an analysis of Margin’s phasing over genic regions to understand its utility for functional studies. With 75× nanopore data from HG001 on GRCh37, we classified each of the GENCODE v35 genes[38] (coding and non-coding) as wholly, partially, or not spanned for the GIAB v3.3.2 high confidence regions, the phasesets proposed by Margin, and the switch errors determined by `whatshap compare` between the two. In Figure 4a, we first plot the number of gene bodies as spanned by high confidence regions (23712 wholly, 26770 partly, 11372 not), then further compare how many of each division were wholly spanned by Margin’s phasesets (22491, 24877, 9807), and finally how many of these had no detected phasing errors (22191, 24563, unknown). In Figure 4b, we plot the number of genes wholly phased by Margin on GRCh38 (60656) for HG003 and HG004 (53817, 55234) and on GRCh37 (62438) for HG001, HG005, HG006, and HG007 (57175, 53150, 53112, 54116).

**Figure 4:**
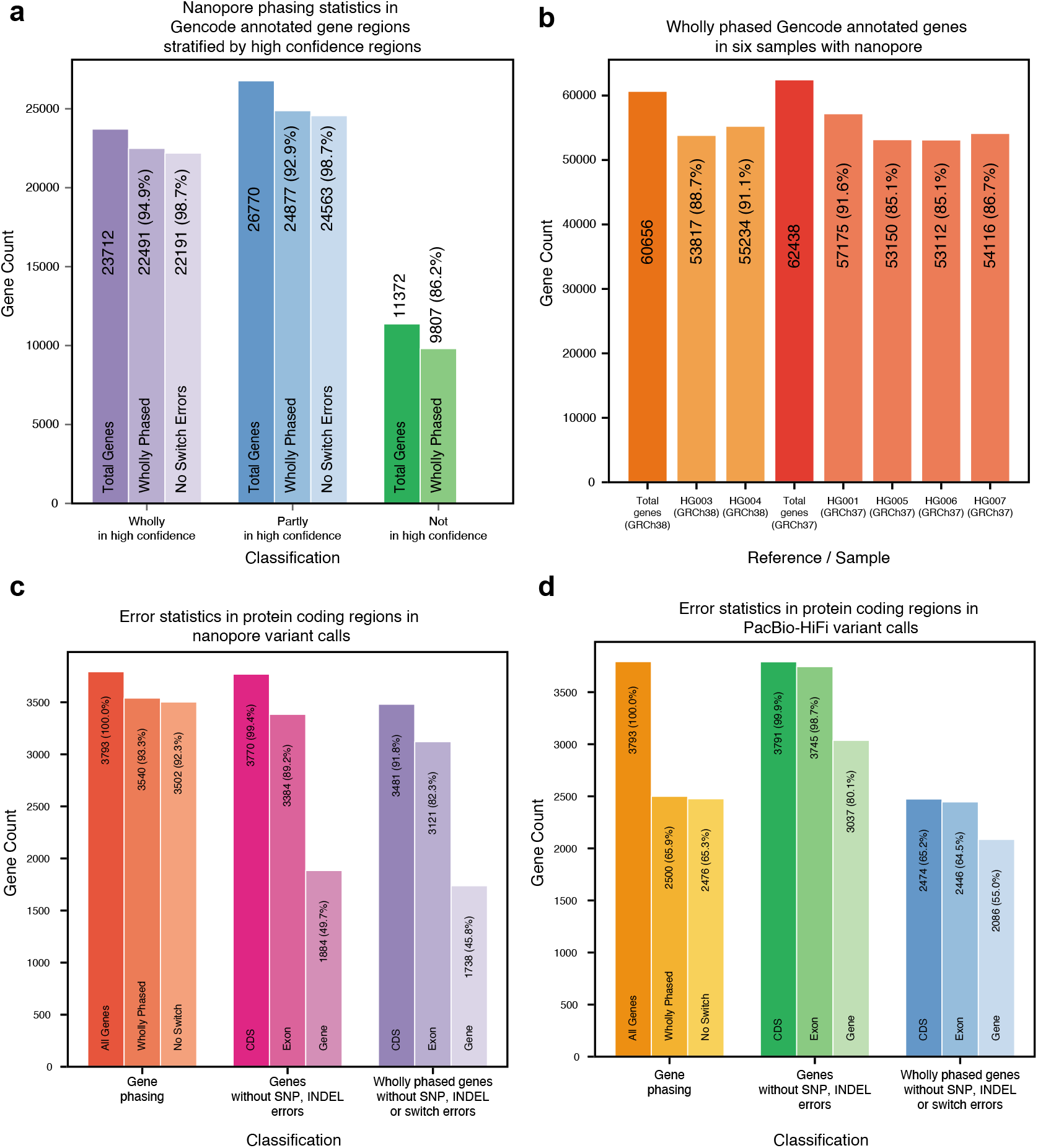
Gene analysis. **(a)** Phasing analysis for HG001 over GENCODE annotated gene regions, stratified by GIAB high confidence coverage. Percentages are relative to their predecessor. **(b)** Wholly phased GENCODE annotated gene regions. Percentages are relative to the total genes annotated on the reference. **(c)** Error statistics including wholly phased genes, genes without SNP or INDEL errors, and wholly phased genes without SNP, INDEL, or switch errors over a subset of the protein coding genes, derived on HG001 from 75x nanopore data. Percentages are relative to total genes in subset. **(d)** The same statistics on HG001 with 35x PacBio HiFi data.

We analyzed accuracy statistics for PEPPER-Margin-DeepVariant with the same HG001 data used above stratified by GENCODE annotations. SNP and INDEL accuracies are largely similar between stratifications of all regions, all genes, and all protein coding genes, with improved performance for protein coding sequence (including CDS, start codon, and stop codon annotations for protein coding genes) (Supplementary Tables 19, 20).

We combined the accuracy and phasing analysis by selecting the 3793 protein coding genes which had at least 80% of their coding sequence covered by the high confidence regions and analyzed the presence of phasing and SNP/INDEL errors on HG001 with 75x nanopore (Figure 4c) and 35x PacBio HiFi (Figure 4d) reads (Supplementary Tables 21, 22). Nanopore had better read phasing for these genes, with 3540 wholly spanned by Margin’s phasesets and only 38 exhibiting a switch error (1.07%), as compared to PacBio HiFi with 2500 wholly spanned genes and 24 switch errors (0.96%). We then counted the number of genes which had no SNP or INDEL errors in the high confidence region for the entire gene, all annotated exons in the gene, and all coding sequences in the gene; PacBio HiFi performs best for this metric with 3037, 3745, and 3791 respectively as compared to nanopore with 1884, 3384, and 3770 perfectly called regions. Lastly, we identified how many of these gene regions were perfectly captured (wholly phased with no switch errors and having no SNP or INDEL miscalls) for the entire gene (1738 for nanopore, 2086 for PacBio HiFi), for all annotated exons (3121, 2446), and for all coding sequences (3481, 2471). For nanopore data, we find that for 91.8% of genes the CDS is fully phased and genotyped without error, and for 82.3% of genes all exons are fully phased and genotyped without error.

### Diploid polishing of *de novo* assemblies

Oxford Nanopore-based assemblers like Flye [16] and Shasta [13] generate haploid assemblies of diploid genomes. By calling and phasing variants against the haploid contigs they produce, it is possible to polish the haploid assembly into a diploid assembly. We implemented such a diploid de novo assembly polishing method with PEPPERMargin-DeepVariant (Figure 5a). It can polish haploid Oxford Nanopore-based assemblies with either Nanopore or PacBio HiFi reads.

**Figure 5:**
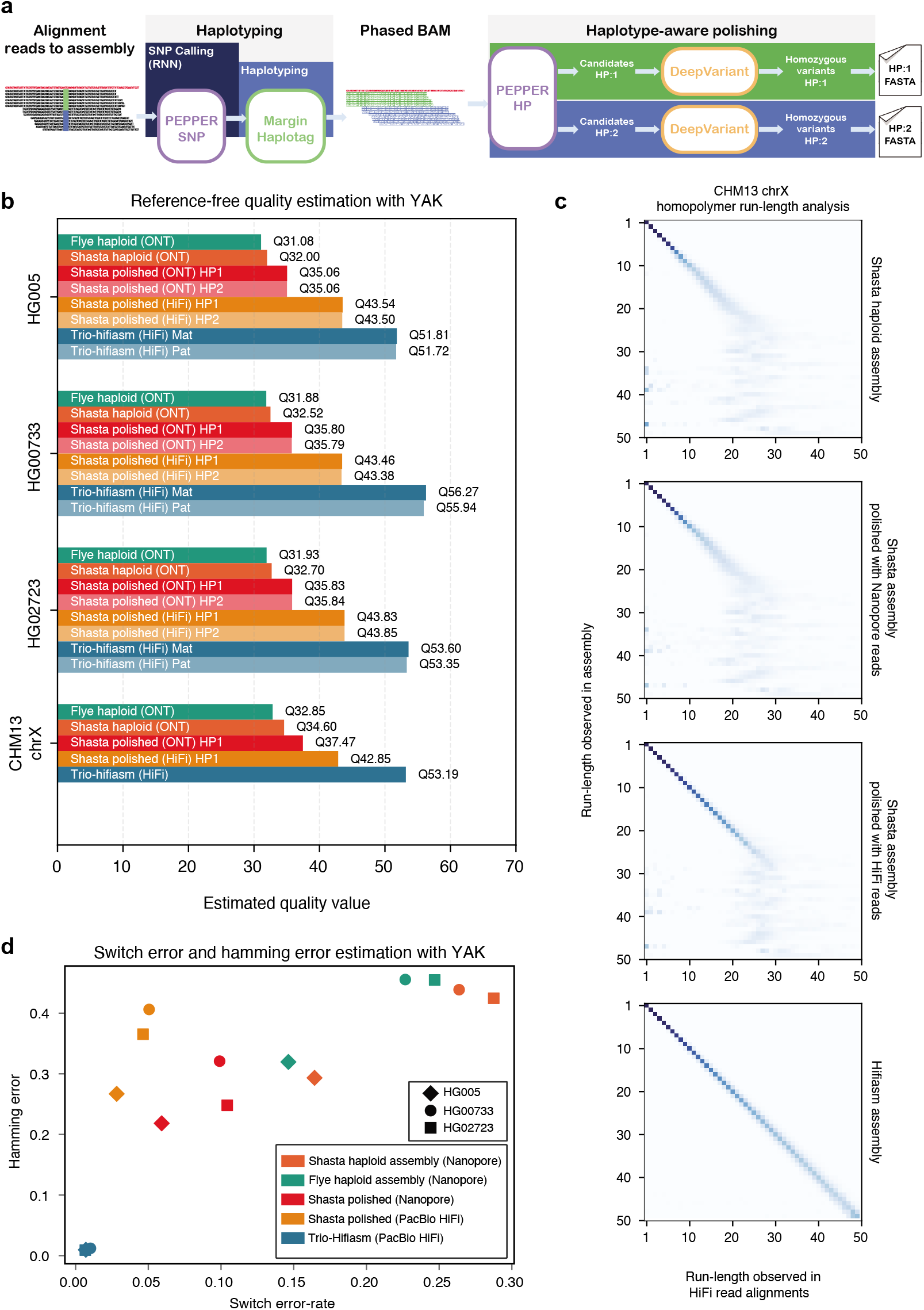
Diploid assembly polishing results. **(a)** Illustration of the diploid assembly polishing pipeline. **(b)** Estimated quality values of assemblies using YAK. **(c)** CHM13-chrX run-length confusion matrix between different assemblies and PacBio HiFi reads aligned to the corresponding assembly. **(d)** Switch error and hamming error comparison between assemblies.

The assembly polishing pipeline employs the modules similarly to the variant calling pipeline. The difference between variant calling and assembly polishing is after we phase the alignment file using the initial set of SNPs, we take candidates from each haplotype independently and classify the candidate as error or not-error using DeepVariant. This entails converting the genotyping classification used in variant calling to a binary classification to predict if a candidate is true error or not. A detailed description of this method is presented in the online methods.

### Diploid *de novo* assembly polishing performance

We generated haploid assemblies using Shasta [13] and Flye [16] for diploid samples HG005, HG00733, HG02723, and haploid sample CHM13 (chrX) using nanopore reads, and we polished the Shasta assemblies using ONT and PacBio HiFi reads. To evaluate the base-level accuracy of the assemblies we use the kmer-based tool YAK[14], which uses Illumina trio data to estimate sequence quality, switch error rates, and hamming error rates. We compare the haploid assemblies, polished diploid assemblies, and trio-aware diploid assemblies generated with hifiasm[14]. Hifiasm uses parental short-read data to generate maternal and paternal assemblies.

The estimated quality values (QV) of nanopore-based assemblies with Shasta (HG005: QV32, HG00733: QV32.7, HG02723: QV32.52) assembler are higher than the nanopore-based Flye assemblies (HG005: QV31.08, HG00733: QV31.93, HG02723: QV31.88). Furthermore, the NG50s of the Shasta assemblies (HG005: 39.83Mbp, HG00733: 42.49Mbp, HG02723: 49.18Mbp) are higher compared to the Flye assemblies (HG005: 37.25Mbp, HG00733: 36.60Mbp, HG02723: 39.65Mbp) (Supplementary table 17).

As Shasta generated higher quality assemblies compared to Flye, we polished the Shasta assemblies with the PEPPER-Margin-DeepVariant diploid polisher. The nanopore-polished assemblies achieve Q35+ estimated quality (HG005: Q35.06, HG00733: QV35.83, HG02723: QV35.8) and PacBio-HiFi-polished assemblies achieve Q40+ estimated quality (HG005: Q43.5, HG00733: QV43.8, HG02723: QV43.8) for all three diploid samples. Finally, we show that the unpolished CHM13-chrX Shasta assembly (QV34.6) can be improved to QV36.9 with nanopore-based and QV42.7 PacBio-HiFi-based assembly polishing with PEPPER-Margin-DeepVariant. Compared to the nanopore-based Shasta assemblies, the trio-aware PacBio HiFi assembler hifiasm achieves higher quality assemblies with respect to base-level accuracy (HG005: QV51.81, HG00733: 53.6, HG02723: 55.94, CHM13-chrX: QV53.03) but the NG50 of the hifiasm assemblies are lower for HG00733 and HG02723 samples (HG005: 51.32Mbp, HG00733: 32.47Mbp, HG02723: 22.21Mbp). In summary, PEPPER-Margin-DeepVariant achieves Q35+ ONT-based assembly polishing and Q40+ PacBio-HiFi-based assembly polishing of ONT assemblies (Figure 5b, Supplementary table 17, Supplementary table 18).

The dominant error modality for ONT data are homopolymers[13]. In Figure 5c we show the run-length confusion matrix of PacBio HiFi read alignments to four chrX assemblies of CHM13-chrX. The Shasta assembly starts to lose resolution at run-lengths greater than 7 (RL-7) and loses all resolution around RL-25. The nanopore-polished assembly improves homopolymer resolution up to RL-10, but also fails to resolve run-lengths greater than RL-25. The PacBio HiFi polished assembly has fair resolution up to RL-25. The trio-hifiasm assembly shows accurate homopolymer resolution up to and beyond RL-50.

Figure 5d shows the switch error-rate of the assemblies. The switch error-rate of haploid Shasta assemblies (HG005: 0.16, HG00733: 0.26, HG02723: 0.28) reduce after polishing with PEPPER-Margin-DeepVariant (HG005: 0.05, HG00733: 0.09, HG02723: 0.10) with ONT data. Similarly, the hamming error rate of the Shasta assemblies (HG005: 0.29, HG00733: 0.43, HG02723: 0.42) reduce after polishing the assemblies with ONT-data (HG005: 0.20, HG00733: 0.31, HG02723: 0.24). Compared to the ONT-polished assemblies the PacBio-HiFi-polished assemblies have higher hamming error-rate (HG005: 0.26, HG00733: 0.40, HG02723: 0.36) but lower switch error-rate (HG005: 0.02, HG00733: 0.04, HG02723: 0.04). The trio-hifiasm that use maternal and paternal short-reads to resolve haplotypes have much lower switch error-rate and hamming error-rate (Figure 5d, Supplementary table 18).

The trio-hifiasm method is able to phase large structural variants in the assemblies. Therefore, trio-hifiasm is expected to produce globally higher quality assemblies. PEPPER-Margin-DeepVariant can not achieve similar global accuracy by polishing haploid assemblies in a diploid manner with small variants. In table 3, we compare HG005 assemblies at the small variant level. The analysis show that the F1-score of unpolished Shasta assembly (INDEL: 0.1203, SNP: 0.4928) improves significantly after polishing with nanopore reads using PEPPER-Marin-DeepVariant (INDEL: 0.3611, SNP: 0.9825). The PacBio-HiFi-polished Shasta assembly achieves similar F1-score (INDEL: 0.9565, SNP: 0.9976) compared to the trio-hifiasm assembly (INDEL: 0.9733, SNP: 0.9988). This analysis provide evidence that PEPPER-Margin-DeepVariant can effectively improve the assembly quality at small variant level.

**Table 3:**
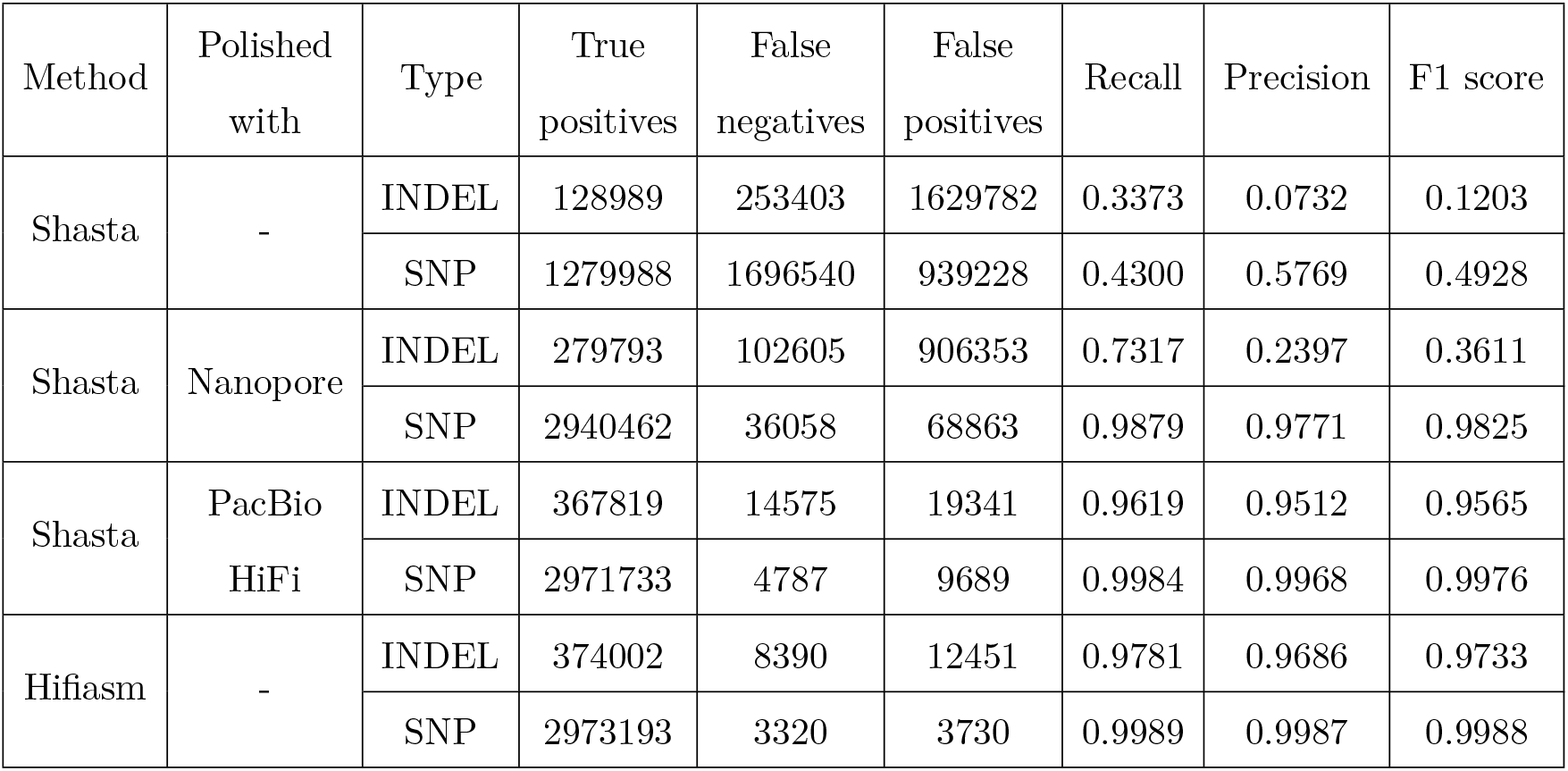
Small variant accuracy evaluation of HG005 assemblies against GIAB HG005 v3.3.2 benchmarking set. We derive a small variant set against GRCh37 from the assemblies using dipcall[39] and compare the variant calls against HG005 GIAB benchmark. We restrict our analysis in regions that are assembled by both Shasta and trio-hifiasm and falls in the high-confidence region defined by GIAB.

## Discussion

Third-generation long-read sequencing technology is allowing gapless human genome assembly [11] and enabling investigations in the most repetitive regions of the genome[28].

In this work, we present PEPPER-Margin-DeepVariant, a state-of-the-art long-read variant calling pipeline for Oxford nanopore data. For the first time, we show that nanopore-based SNV identification outperforms a state-of-the-art short-read based method at whole genome scale. Particularly in segmental duplication and difficult-to-map regions, the nanopore-based method outshines the short-read based method. It seems likely, therefore, that the anticipated widespread application of long-read variant calling will for the first time accurately illuminate variation in these previously inaccessible regions of the genome.

The genomic contexts where nanopore SNP accuracy suffers for our pipeline are identifiable, meaning that variant calls in these regions can be treated with skepticism while calls outside these contexts can be handled with confidence. The one obvious area that Nanopore variant calling lags is in INDEL accuracy. While the results achieved here are to our knowledge the best shown so far, we believe it is likely that further technological innovations at the platform level will be required to make nanopore INDEL accuracy on par with other technologies.

Oxford Nanopore provides a highly-multiplexed sequencing solution with its PromethION device [13]. With this device and the PEPPER-Margin-DeepVariant pipeline described here it should be comfortably possible to go from biosample collection to complete genome inferences in under half a day. This fast turnaround should enable its use in a medical context, where diagnosis for acute disease situations requires speed.

We have demonstrated our nanopore-based phasing is able to wholly phase 85% of all genes with only 1.3% exhibiting a switch error. This phasing ability could play a useful role in population genetics studies [40, 41] and clinical genomics [42]. For clinical applications the accurate identification of compound-heterozygotes should be particularly valuable.

We have extended PEPPER-Margin-DeepVariant to PacBio HiFi reads and demonstrated a more accurate and cheaper solution to the existing WhatsHap-DeepVariant variant calling methods, making cohort-wide variant calling and phasing with PacBio HiFi more accessible.

We have demonstrated diploid polishing of nanopore-based haploid assemblies with PEPPER-Margin-DeepVariant. We achieve Q35+ nanopore polished assemblies and Q40+ PacBio-HiFi-polished assemblies. We observe that our polishing method can resolve homopolymer errors up to 20bp with PacBio HiFi data. However, our polishing method fails to resolve 25bp+ long homopolymers indicating that they need to be resolved during the consensus generation of the *de novo* assembly methods. As nanopore assembly methods like Shasta move toward generating fully resolved diploid genome assemblies like trio-hifiasm, our polishing method can enable nanopore-only Q40+ polished diploid assemblies.

## Supporting information

Supplementary Results

## Acknowledgements

Research reported in this publication was supported by the National Human Genome Research Institute of the National Institutes of Health under Award Numbers U41HG010972, 1R01HG008742, U01HG010961, R01HG009737 and 2U41HG007234. Research reported in this publication was supported by the National Heart, Lung, And Blood Institute of the National Institutes of Health under Award Number U01HL137183. We also acknowledge Circulomics Inc. for sharing HG001 Nanopore data. The content is solely the responsibility of the authors and does not necessarily represent the official views of the National Institutes of Health.

## Code Availability

The modules of PEPPER-Margin-DeepVariant are publicly available in these repositories:

- PEPPER: https://github.com/kishwarshafin/pepper
- Margin: https://github.com/UCSC-nanopore-cgl/margin
- DeepVariant: https://github.com/google/deepvariant

For simpler use we have also created a publicly available docker container `kishwars/pepper_deepvariant:r0.4` that can run our variant calling and polishing pipelines.

## Data availability

### Analysis data available

We have made the analysis data available publicly (variant calling outputs, genome assemblies etc.) in: https://console.cloud.google.com/storage/browser/pepper-deepvariant-public/analysis_data.

### Sequencing data availability

We used several publicly available datasets:

- Genome-In-a-Bottle consortium[26, 28]: ftp://ftp-trace.ncbi.nlm.nih.gov/giab/ftp/data/
- Human Pangenome Reference Consortium (HPRC): https://s3-us-west-2.amazonaws.com/human-pangenomics/index.html
- Telomere-to-telomere consortium[11, 12]: https://github.com/nanopore-wgs-consortium/CHM13

Please see supplementary notes to find specific links to the sequencing data that we used for our analysis.

## Online Methods

### Analysis methods and data pre-processing

#### Read alignment

We used `minimap2` [43] version `2.17-r941` and `pbmm2` version `1.4.0` to align reads to a reference genome. The supplementary notes have details on execution parameters.

#### Subsampling alignment files to different coverages

We used `samtools` [44] version `1.10` to generate alignment files of different coverages. The supplementary notes have details on execution parameters.

#### Variant calling

We used the following methods to call variants with nanopore data:

- `PEPPER-Margin-DeepVariant` version `r0.4`.
- `Medaka` [18] version `v1.2.1`.
- `Clair` [19] version `v2.1.1`.
- `Longshot` [20] version `v0.4.2`.

For Illumina short-reads and PacBio HiFi we used `DeepVariant` version `v1.1.0`. The details on execution parameters are available in supplementary notes.

#### Benchmarking variant calls

We used `hap.py` [45] version `v0.3.12` to assess the variant calls against GIAB truth set. The `hap.py` program is available via `jmcdani20/hap.py:v0.3.12` docker image. The command used for the assessment is described in the supplementary note.

For HG002, HG003, HG004 we used GIAB v4.2.1 truth set against `GRCh38` reference and for HG001, HG005, HG006, HG007 samples, we used v3.3.2 variant benchmarks against `GRCh37` reference genome. We used GIAB stratification `v2.0` files with `hap.py` to derive stratified variant calling results. The GIAB benchmarking data availability is listed in data availability section of supplementary notes.

#### Phasing and haplotagging

We used `Margin` version `v2.0` and `WhatsHap` [22] version `v1.0` to haplotag and phase the variants. `Margin` is available in https://github.com/UCSC-nanopore-cgl/margin and `WhatsHap` is available in https://github.com/whatshap/whatshap. The details on how we ran these tools is described in supplementary notes.

#### Small variant switch error rate and hamming error rate determination

We used a Workflow Description Language (WDL)-based analysis pipeline `whatshap.wdl` available in https://github.com/tpesout/genomics_scripts to derive the switch error rate and hamming error rate compared to the GIAB truth set. The `whatshap.wdl` workflow invokes the `stats` and compare submodules available in `whatshap` version `v1.0`.

In our analysis, we compared phased variants against GRCh37 reference against GIAB v3.3.2 truth set to derive switch error rate and Hamming error rate. We only considered variants that have `PATMAT` annotation in the truth set and that fall in the high-confidence region defined by GIAB benchmarking set. The non-`PATMAT` annotated variants in the GIAB benchmarking variant set are not trio-confirmed so we did not use those to benchmarking our phasing methods. We used `whatshap compare` command to generate the whole genome switch error rate and used a custom script defined in `whatshap.wdl` to derive the hamming error rates.

#### Local phasing correctness calculation

For the Local Phasing Correctness (LPC) analysis, we used the `calcLocalPhasingCorrectness` executable found in the https://github.com/UCSC-nanopore-cgl/margin repository. The LPC analysis require a truth variant set and a query variant set. We used GIAB benchmarking set as truth. The `calcLocalPhasingCorrectness` generates a `tsv` file describing the results.

We used https://github.com/tpesout/genomics_scripts/plot_haplotagging_lpc.py script to visualize the results. The details of parameters is described in supplementary notes. The methods used in local phasing correctness as a metric is presented separately in the methods description.

#### Haplotagging accuracy and natural switch determination

We used https://github.com/tpesout/genomics_scripts/haplotagging_stats.py to calculate the hap-lotagging accuracy. The script calculates average haplotagging accuracy and average tagged reads per 10kb. Details on execution parameters is available in supplementary notes.

We visualized the natural switch error using `compare_read_phasing_hapBam.py` available in https://github.com/tpesout/genomics_scripts. Details on execution parameters is available in supplementary notes.

#### Phaseblock N50 calculation

An N50 value is a weighted median; it is the length of the sequence in a set for which all sequences of that length or greater sum to 50% of the set’s total size. We used the `ngx_plot.py` available from https://github.com/rlorigro/nanopore_assembly_and_polishing_assessment/ to plot phaseblock N50. From a phased VCF file, we extracted the phaseblock name, contig, start position, and end position to create the input file. We used `3272116950` as the size of the genome to maintain consistency with previous work [13].

#### Variant calling and phasing analysis on Gencode annotated regions

We used Gencode v35 [37] to determine the variant calling and phasing accuracy in gene regions. The Gencode data is publicly available and can be found in data availably section of supplementary notes. We used https://github.com/tpesout/genomics_scripts/gencode_to_stratification.py script to convert the Gencode regions to a bed file that is acceptable to `hap.py`. With the newly defined stratified regions from Gencode, we ran `hap.py` to determine the variant calling accuracy in gene regions.

#### Nanopore and PacBio HiFi *de novo* assembly generation

We used `Shasta` [13] with a development build after version `0.7.0` available from https://github.com/chanzuckerberg/shasta (commit `06a639d36d26a4203c0b934d6e63c719750c5398`) and `Flye` [16] version `2.8.2` available from https://github.com/fenderglass/Flye to generate nanopore-based *de novo* assemblies. For PacBio-HiFi-based assemblies we used `hifiasm` [14] version `0.14` available from https://github.com/chhylp123/hifiasm. The commands used to generate the assemblies are provided in execution parameters section of supplementary notes.

#### Assembly QV and switch error rate analysis

We assessed the assemblies with `yak` [14] version 0.1 available from https://github.com/lh3/yak. YAK is a short-read kmer-based assembly quality estimator. We use short-reads for each sample to estimate the quality of the assembly with *k-mer* size of 31, With parental short-reads, YAK can also estimate the switch error rate in the assembly. WDL version of the pipeline `standard_qc_haploid.wdl` is available in https://github.com/human-pangenomics/hpp_production_workflows/.

#### Homopolymer run-length analysis

We used `runLengthMatrix` module of `margin` to derive the homopolymer run-length analysis between assembly and reads. In `runLengthMatrix`, we convert each read sequence into RLE form and track a map of raw positions to RLE positions. We convert from a raw alignment to RLE alignment by iterating through the matches in raw space and tracking the previous RLE match indices. From this set of matched read and reference RLE positions, we construct a confusion matrix. Details of the command are provided in execution parameters section of supplementary notes.

#### Small variant accuracy evaluation of assemblies

We used `dipcall` [39] to identify the small variants from the assemblies. The `dipcall` variant identification takes the maternal and paternal haplotypes generated by a phased assembly and a reference genome sequence. It maps the haplotypes to the reference and generates a VCF file containing all small variants identified in the assembly. For the haploid assembly, we provided the haploid assembly as both maternal and paternal haplotypes to `dipcall. dipcall` also generates a bed file containing regions where the assembly maps to the reference. We intersected the bed files to get regions that are assembled by all assembly methods and intersected with GIAB high-confidence region. Finally, we used `hap.py` to compare the variant calls derived from the assemblies against GIAB benchmarking VCF to get the accuracy statistics. Please see supplementary notes for `dipcall` parameters.

### Method description

#### PEPPER

PEPPER is a recurrent neural network-based sequence prediction tool. In PEPPER, we use summary statistics derived from reads aligned to a reference to produce base probabilities for each genomic location using a neural network. We translate the position-specific base probabilities to the likelihood values of candidate variants observed from the read alignments. We propose candidate variants with likelihood value above a set threshold to DeepVariant for genotyping. Candidate pre-filtering with PEPPER ensures a balanced classification problem for DeepVariant and achieves high-quality variant calling from erroneous long-reads.

We use PEPPER in two steps in the variant calling pipeline. Initially, we use the PEPPER-SNP submodule to find single nucleotide variants (SNVs) from the initial unphased alignment file. In this setup, we tune PEPPER-SNP to have high precision so Margin can use the SNVs confidently to phase the genome. To this end, we also exclude INDELs from the callset as they have notably worse performance for nanopore reads. Margin can then tag reads in the alignment file with predicted haplotypes.

After Margin, we use the PEPPER-HP submodule on the phased alignment file to generate haplotype-specific likelihoods for each candidate variant observed from the read alignments. In PEPPER-HP, we consider SNVs, insertions, and deletions (INDELs) as potential candidate variants. We propose the candidate variants with likelihood values higher than a set threshold to DeepVariant for genotyping with a more extensive convolutional neural network (CNN). We tune PEPPER-HP to achieve high-sensitivity but low-precision during candidate finding. The PEPPER-Margin-DeepVariant suite can identify small variants with high-quality from erroneous reads.

#### PEPPER-SNP

PEPPER-SNP is a submodule of PEPPER used to identify single nucleotide variants (SNVs) from reads aligned to a reference sequence. PEPPER-SNP works in three steps: image generation, inference, and candidate finding. First, we generate summary statistics from reads aligned to a reference sequence. We encode basic alignment statistics at each genomic location in an image-like tensor format. Second, we apply a recurrent neural network to predict the two most likely bases at each genomic location. Finally, we use the base predictions from each genomic location to compute the likelihoods of SNVs we observe from the reads. We filter candidate variants with a likelihood value below a set threshold to find a set of SNVs. The SNV set we get from PEPPER-SNP is used by Margin to phase the alignment file.

#### PEPPER-SNP: Image generation

In the image generation step of PEPPER-SNP, we generate summary statistics of base-level information per genomic location. The summary provides weighted observation of bases from all reads divided into nucleotide and orientation.

In PEPPER-SNP, we do not encode insert bases observed in reads to reference alignment as we only look for SNVs. We use a position value to represent a location in the reference sequence. For each genomic location, we iterate over all reads that align to that genomic location and encode ten features to encode base-level information: {*A, C, G, T, Gap*(*)} divided into two read orientations: {*forward, reverse}*. Finally, we normalize the weights of each genomic position based on the read coverage.

Figure 6 describes the feature encoding scheme we use in the image generation step of PEPPER-SNP. The top row of the image, annotated as *REF*, describes the reference base observed at each genomic location. The colors describing the bases are {*A: Blue,C: Red,G: Green,T: Yellow,Gap*(*): *White}*. Each row after *REF* describes a feature; each feature encodes an observation of nucleotide bases from a forward, or a reverse strand read. We use ten features to encode base-level information: {*A,C,G,T,Gap*(*)} divided into two read orientations: {*forward, reverse*}. For example, *A_F_* encodes the observations of base *A* from forward-strand reads, and *A_R_* encodes observations of base *A* from reverse-strand reads. The columns describe genomic locations to the reference sequence.

**Figure 6:**
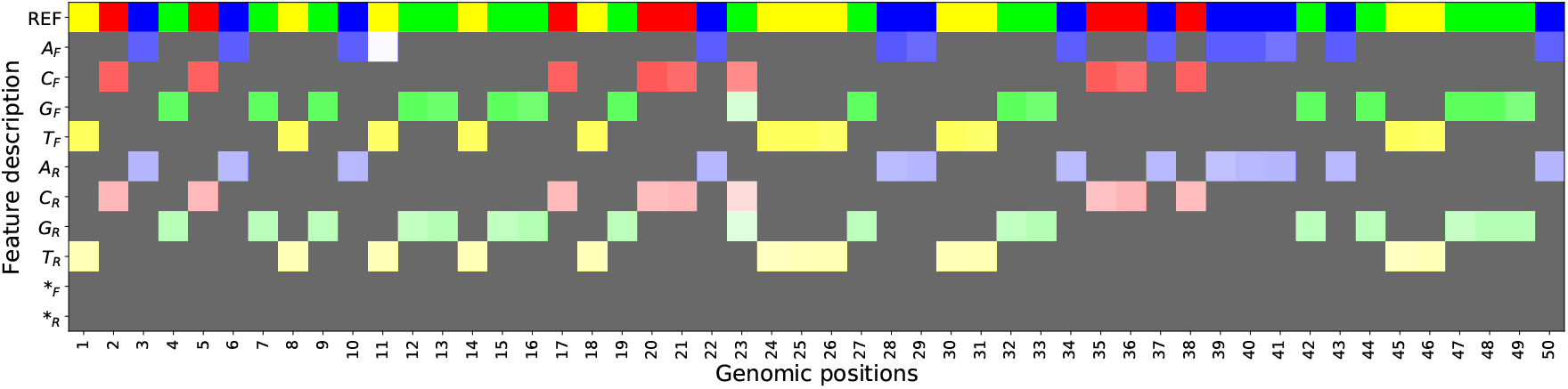
PEPPER-SNP image generation scheme.

In each column, we encode each observation as weights, which we show as alpha of each base. The grey weights are zero weights. At position (23), the weight distribution of *{C, G*} bases indicates a potential heterozygous variant at that position. The *REF* row is shown in the figure to describe the scheme; in practice, we do not encode the *REF* row.

#### PEPPER-SNP: Inference model

In the inference step of PEPPER-SNP, we use a recurrent neural network for sequence prediction. The network architecture consists of two bidirectional gated recurrent unit (GRU) layers and a linear transformation layer. The linear transformation layer produces a prediction of two bases for each genomic location present in the summary image. To identify potential variants, we use 15 class-labels for base prediction: {*AA, AC, AG, AT, A*, CC, CG, CT, C*, GG, GT, G*, TT, T*, **}*. We do not use co-linear classes like *CA* and *AC* as two separate classes, as it is not possible to differentiate between these two classes from the summary observations.

From the image-generation module of PEPPER-SNP, we get summary images in 1kb chunks. We use a sliding window method during inference and chunk the 1kb segments into multiple overlapping windows of 100bp segments. We first run inference on the leftmost window and go to the next window with 50bp overlapping bases. We pass the hidden state output from the left window to the next window and keep a global inference counter to record base-predictions.

Figure 7 describes the neural network-based inference scheme. The two dotted boxes indicate two adjacent windows with overlapping sequences. The top panel shows the inference scheme. We start with the first window and produce base predictions for each genomic location present in that window; then, we slide the window to the right. We do this from left to right on the entire genomic sequence. We record the base predictions from all windows in a global counter and report them to the candidate finder to calculate candidate likelihoods.

**Figure 7:**
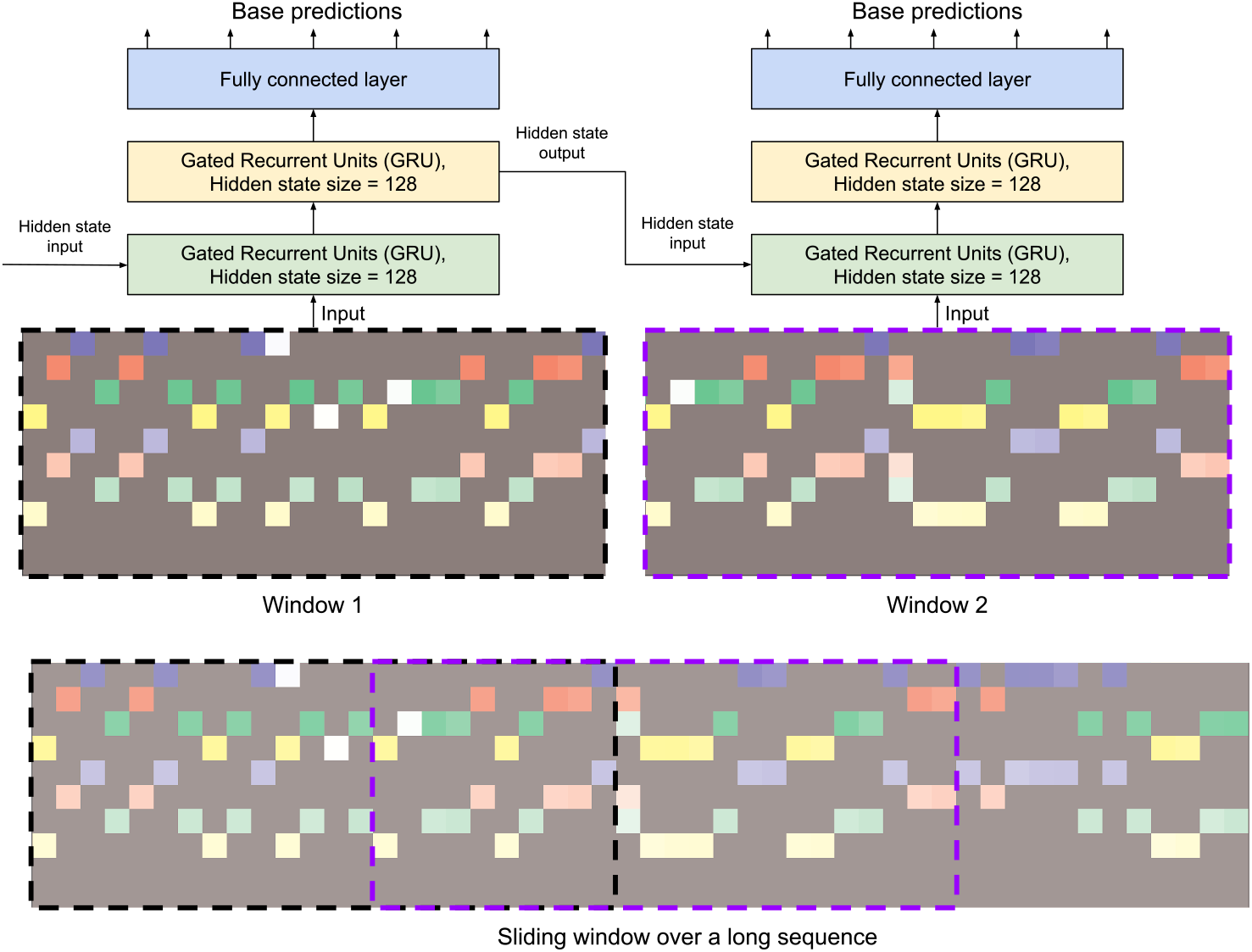
PEPPER-SNP inference scheme.

We trained the inference model using a gradient descent method. We use adaptive moment estimation (Adam [46]) to compute gradients based on a cross-entropy loss function. The loss function is defined to calculate the prediction performance at each genomic location against a labeled set of expected base observations derived from the Genome-In-A-Bottle (GIAB) truth set. The gradient optimization attempts to minimize the loss function by tuning the parameters of the neural network.

We trained PEPPER-SNP with 100 × coverage of HG002 data subsampled at different coverage values. We split the training sets into three sets: train, test, and holdout. We use chromosome 1 to 19 for training, chromosome 20 for testing, and we keep 21 and 22 as holdout sets. We train the models for several epochs and test after each epoch. Finally, we pick a model that performs the best on the holdout dataset.

#### PEPPER-SNP: Candidate finding

In the candidate finding step of PEPPER-SNP, we take the base-predictions and derive likelihoods of SNVs we observe from the read alignments. If the likelihood value of a variant is above a set threshold, we pick that allele to be a real variant.

In PEPPER-SNP, we derive a allele probability (*AP*) and a non-reference observation likelihood (*NR*) for each observed SNV. First, at each genomic location (*pos*) we use the prediction vector of base-classes {*AA, AC, AG, AT, A*\*, *CC, CG, CT,C*\*, *GG, GT, G*\*, *TT, T*\*, **} to derive two prediction vectors *V*_1_ [*pos*] = [*A_1_,C_1_,G_1_,T_1_,Gap(*)_1_*] and *V*_2_[*pos*] = [*A*_2_,*C*_2_,*G*_2_,*T*_2_,*Gap*(*)2]. For example, the predicted value of classs {*GT*} contributes to *V*_1_[*pos*][*G*_1_] and *V*_2_[*pos*][*T*_2_] values. Then, we iterate over all the reads to find potential SNVs by recording each read base that does not match the reference base. We calculate the likelihood of candidate base *b*, observed at position *pos*, by taking the maximum likelihood from the prediction vectors *V*_1_ [*pos*] and *V*_2_ [*pos*] as shown in equation 1 where we denote allele likelihood as *AP*.

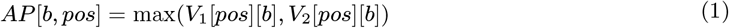

We also calculate the likelihood of non-reference base observation (*NR*) to estimate the likelihood of observing any allele other than the reference allele at a location. We derive non-reference base observation likelihood from prediction vectors *V*_1_ and *V*_2_ independently and take the maximum value between two values. For each prediction vector, we take the sum of the values, subtract the observation likelihood of the reference base, and divide by the sum of the prediction vector. Equation 2 describes the calculation *NR* in PEPPER-SNP.

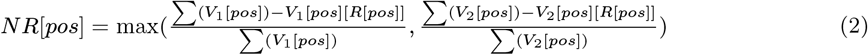

Finally, we derive a genotype for the variant from the prediction vector *V*_1_ and *V*_2_. If a variant has a likelihood above a set threshold observed in both *V*_1_ and *V*_2_, we set the genotype as a homozygous alternate (1/1). If the likelihood is above the threshold in one vector but not in the other, we call it a heterozygous variant (0/1). For each variant, we use *NR* value of that position to be the genotyping quality.

#### PEPPER-SNP: Code availability

PEPPER-SNP is available at https://github.com/kishwarshafin/pepper.

#### PEPPER-HP: Haplotype-aware sequence prediction

PEPPER-HP is a haplotype-aware sequence prediction tool designed to find candidate variants from read alignments. In PEPPER-HP, we take a phased alignment file as input where each read has a haplotag of {0,1, 2} indicating which haplotype the read represents (or lack of haplotype information), and we output a set of SNV, insertion, and deletion candidates for genotyping using DeepVariant.

Similar to the PEPPER-SNP submodule, PEPPER-HP has three steps, image generation, inference, and candidate finding. In the image generation step, we generate two sets of summary statistics, one per haplotype, and save them as image-like tensors. We use a recurrent neural network to predict bases on each haplotype for each genomic location in the inference step. Finally, we calculate likelihood values for SNV and INDEL candidates based on base-predictions of each haplotype. We consider candidates with likelihood values over a certain threshold to be candidate variants and propose them to DeepVariant for genotyping.

#### PEPPER-HP: Image generation

In the image generation step of PEPPER-HP, the input is an alignment file with phased reads, and we generate image-like summary statistics of base-level information per genomic location for each haplotype {0, 1}. The summary of haplotype 1 provides weighted observation of bases from reads with haplotag 1 divided into nucleotide and orientation. Similarly, the haplotype 2 summary provides weighted observations of bases from reads with haplotag 2. Reads that are unphased or have haplotag 0 contribute to summary statistics for both haplotypes.

In PEPPER-HP, we represent position in reference sequence using two values: position and index. The position value indicates a location in the reference sequence, and we use the index to accommodate insert alleles anchored to a position. All reference sequence positions have an index of 0. On each haplotype {0, 1}, we iterate over all haplotype associated reads that align a genomic location and encode ten features to encode base-level information: {*A,C,G,T,Gap*(*)} divided into two read orientations: {*forward,reverse*}. The weights depend on the mapping quality and base quality of the associated reads. Finally, we normalize the weights of each genomic position based on the haplotype associated read coverage.

In Figure 8, we describe the feature encoding scheme of PEPPER-HP. We derive two summary statistics based on the haplotype association of the reads. The top row of the image, annotated as *REF*, describes the reference base observed at each genomic location. The colors describing the bases are {*A: Blue, C: Red,G: Green, T: Yellow, Gap*(*): *White*}. Each column represents a reference position with two values (*pos, index*). For example, pos (14, 0) is the reference reference position 14 and (14,1) is the insert base anchored in position (14, 0). For each haplotype, we use ten features to encode base-level information: {*A,C,G,T,Gap*(*)} divided into two read orientations: *{forward, reverse*}. From the summary, we can see that at location (23, 0) HP-1 observers *C* bases where the reference is *T* but in HP-2 the reads observe *C* bases that match with the reference, denoting a heterozygous variant present in haplotype-1 sequence.

**Figure 8:**
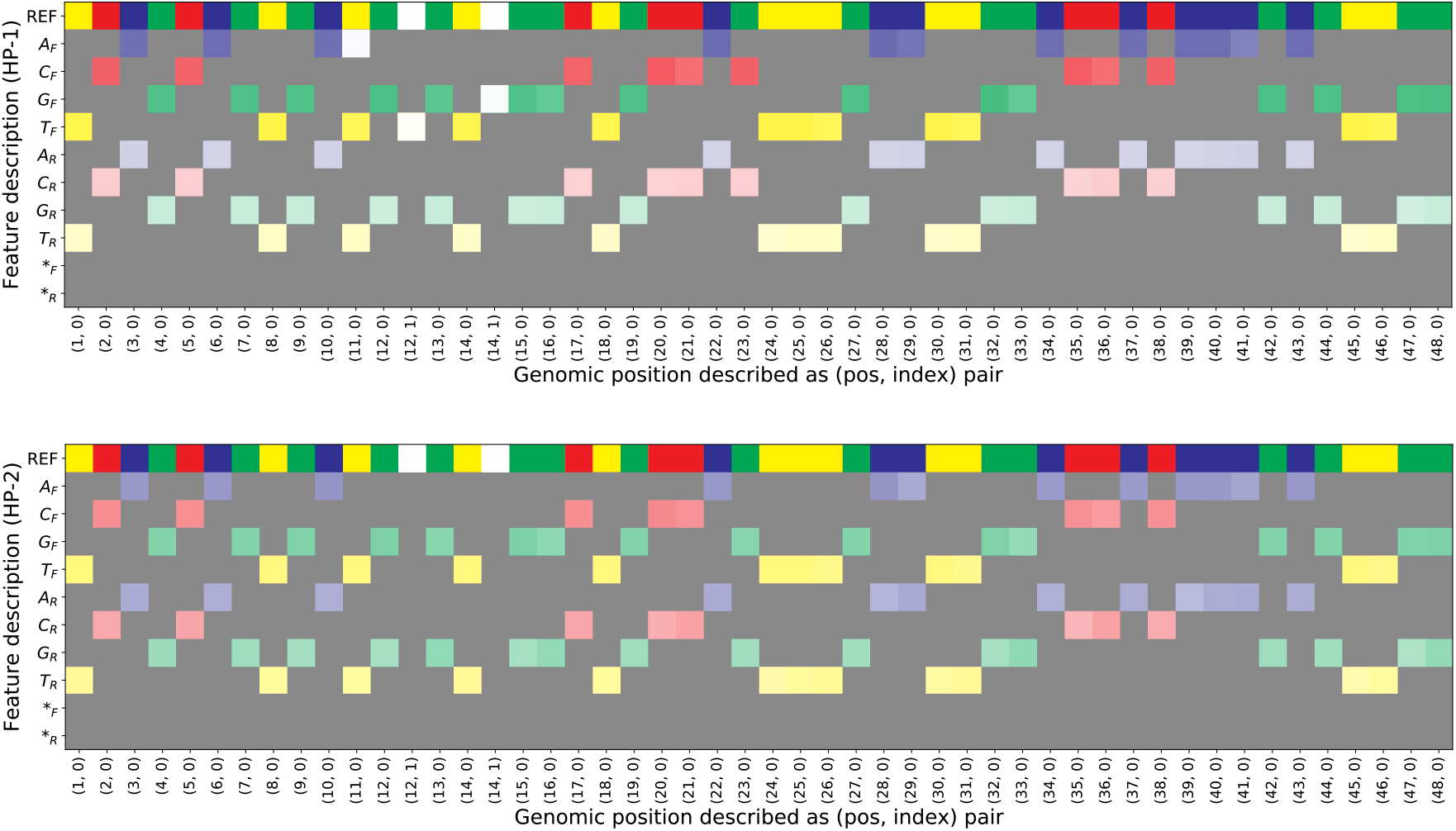
PEPPER-HP haplotype specific image generation scheme. Each row describes an encoded feature and each column describes a reference position. The top summary is derived from reads with haplotag 1 (HP-1) and the bottom is derived from reads with haplotag 2 (HP-2).

#### PEPPER-HP: Inference model

We use a recurrent neural network for sequence prediction on haplotype-specific images of PEPPER-HP. The network architecture consists of two bidirectional gated recurrent unit (GRU) layers and a linear transformation layer. For each haplotype, the linear transformation layer predicts a base for each genomic location present in the image-like tensor. We use five class-labels for base prediction: {*A, C, G, T, Gap*(*)}.

The haplotype-specific images in 1kb chunks, and we use a sliding window method to go over the 1kb chunk in overlapping 100bp segments on both haplotype images simultaneously. We start from the leftmost window of 100bp and slide the window to 50bp to the right for the next step. We use two global counters from the base predictions, one per haplotype, to record the haplotype-specific base predictions.

The inference scheme of PEPPER-HP is shown in Figure 9. We have two haplotype-specific image for each genomic region representing two haplotypes. For each haplotype, we start from the leftmost window and generate haplotype-specific base predictions. The base predictions are recorded in two global counters.

**Figure 9:**
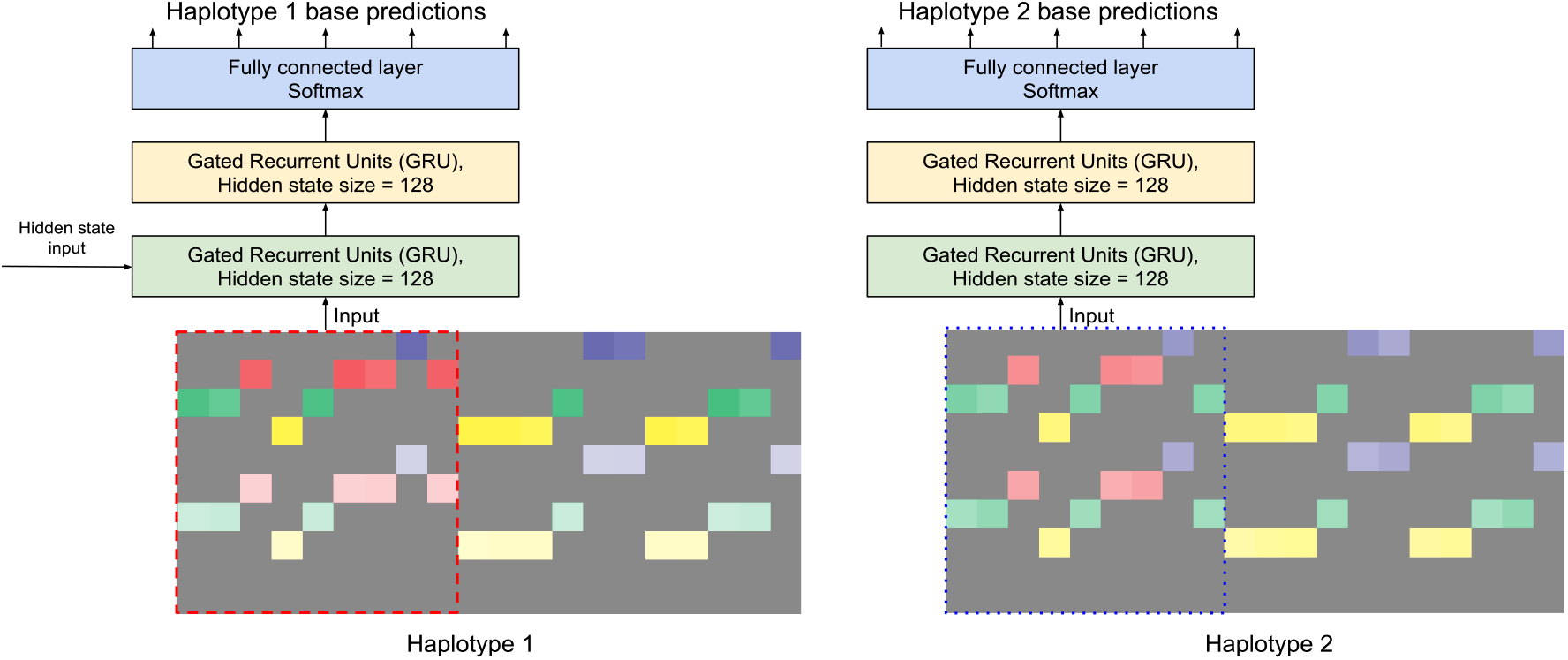
PEPPER-HP haplotype-specific inference scheme.

We train the PEPPER-HP inference model using a gradient descent method. We use adaptive moment estimation (Adam) to compute gradients based on a cross-entropy loss function. The loss function is defined to calculate the prediction performance at each genomic location against a labeled set of expected base observations derived from the Genome-In-A-Bottle (GIAB) truth set. We use GIAB v3.3.2 truth set as the variants in v3.3.2 are phased. We use phase-specific base predictions to optimize PEPPER-HP model for each haplotype.

We train PEPPER-HP with three sets of HG002 data with 50×, 80 ×, and 100 × coverage. We further generate multiple train sets by arbitrarily downsampling the three training sets at different fractions. We split the training sets into three sets: train, test, and holdout. We use chromosome 1 to 19 for training, chromosome 20 for testing, and we keep 21 and 22 as holdout sets. We train the models for several epochs and test after each epoch. Finally, we pick the model that performs the best on the holdout dataset.

#### PEPPER-HP: Candidate finding

In the candidate finding step of PEPPER-HP, we evaluate variants observed from the read alignments. We use the base-predictions from the neural network to calculate the likelihood of an observed allele. If the likelihood value of a candidate variant is above a set threshold, we pick that variant as a potential candidate for DeepVariant to assess.

We evaluate the SNVs and INDELs observed in read alignments to find potential candidate variants. First, at each genomic location (*pos, index*) we use the haplotype-specific prediction values of base-classes *V*_H1_ [*pos, index*] = [*A*_1_,*C*_1_,*G*_1_,*T*_1_,*Gap*(*)_1_] and *V*_H2_[*pos, index*] = [*A*_2_, *C*_2_, *G*_2_, *T*_2_, *Gap*(*)_2_]. We iterate over all the reads to find potential variants by recording the read base that does not match the reference base. We calculate the allele likelihood of a SNV candidate *AP_SNV_* with base observation *b*, observed at position *(pos, index),* by taking the maximum likelihood from the prediction vectors *V*_H1_[(*pos, index*)] and *V*_H2_[(*pos, index*)] as described in equation 3. For inserts and deletes we extend the likelihood calculation to cover the length of the allele.

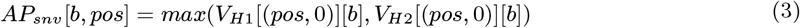

We also calculate the likelihood of observing a variant other than the reference allele at any location. In equation 4, let *R*[(*pos, index*)] be the reference base at location (*pos, index*) and *N* be the maximum index value observed in position *pos*. The reference base at any position with *index* > 0 is *gap*(*). We take the maximum value of observing a non-reference base between (*pos*, 0) and (*pos, max_index*[*pos*]). For each index, we calculate the total value of the prediction vector, subtract the observation likelihood of the reference base and divide by the sum of the prediction vector. The non-reference observation likelihood *NR* is the maximum value we observe across index values. For insertion and deletion alleles we cover the allele length and take the maximum value as the *NR* for those candidates.

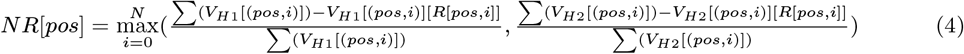

Based on the allele likelihood and non-reference observation likelihood, we calculate a likelihood value for each type of candidate SNP, insert and delete. Then for each type, we set a threshold value and if a candidate passes the threshold value, we propose the candidate to DeepVariant for genotyping.

#### PEPPER-HP: Code availability

PEPPER-HP is available at https://github.com/kishwarshafin/pepper as a submodule.

#### Margin

Margin is a suite of tools employing Hidden Markov Models (HMMs) to perform genomic analysis with long reads. MarginPhase (the first module) was introduced alongside WhatsHap as a tool performing joint genotyping and phasing [22]. MarginPolish (the second module) was introduced as a graph-based assembly polisher which can do standalone polishing and is the first step in a two-part polishing framework MarginPolish-HELEN [13]. Release 2.0 of Margin incorporates both tools into one suite, including diploid-aware polishing in MarginPolish which has informed improvements in a new iteration of MarginPhase.

In this paper we focus exclusively on improvements made to the phasing submodule. The core partitioning algorithm is described in our previous work [22], but we provide a summary here of the previous methodology, followed by a description of the modifications made in the current iteration. First, we give a high-level overview of the phasing workflow.

#### Overview

Margin takes as input an alignment (BAM), reference (FASTA), and variant set (VCF). It determines regions to run on from the alignment and variant set, and it breaks the input into chunks to enable multiprocessing. For each chunk, it extracts reference substrings around each variant site, and read substrings aligning around each variant site. For each variant site, it calculates the alignment likelihood between all substrings and all variant alleles. These likelihoods are used in the core phasing algorithm which bipartitions reads and assigns alleles to haplotypes. After all chunks have been analyzed, we stitch the chunks together to produce results across whole contigs. Last, we output a copy of the input BAM with haplotagged reads and a copy of the input VCF with phased variants.

##### Parameterization

Margin is parameterized with a configuration file. For each configuration, parameters are grouped into `polish` and `phase` sections. There is overlap in the parameters used by both submodules, so some parameters used by the phasing algorithm fall under the `polish` heading and vice versa. When referenced in this document, we specify the full path of the parameter and the default value associated with the intended configuration.

#### Core Phasing Algorithm

For the core phasing algorithm, we construct a graph *G* = (*V_G_, E_G_*) describing all possible bipartitions of reads with positions as variant sites, vertices as a combination of position and read bipartitions, and edges as possible transitions between bipartitions for adjacent positions. For example, at postition *P_i_* with aligned reads *R*_1_ and *R*_2_, we have the possible vertices *V*_*i*_0__ with haplotypes *H*_1_ = {*R*_1_,*R*_2_} and *H*_2_ = {} (*R*_1_, *R*_2_), *V*_*i*_1__ = *R*_1_/*R*_2_, *V*_*i*_2__ = *R*_2_/*R*_1_, and *V*_*i*_3__ =./*R*_1_ *R*_2_. At position *P_j_* with the same aligned reads *R*_1_, *R*_2_ and a new read *R*_3_, vertex *V*_*j*_0__ = *R*_1_, *R*_2_, *R*_3_/. and *V*_*j*_1__ = *R*_1_, *R*_2_/*R*_3_ are both connected to vertex *V*_*i*_0__ because all reads shared between vertices are have the same haplotype assignment, but *V*_*j*_2__ = *R*_1_/*R*_2_, *R*_3_ is not connected to *V*_*i*_0__ because read *R*_2_ has different haplotype assignments in the two vertices. We extend each vertex as described above to additionally represent all possible genotypes. After running the forward-backward algorithm on this graph, at each position the posterior distribution over states describing read bipartitions and genotypes can be marginalized to determine the most likely genotype.

The state space for this algorithm increases exponentially with the number of reads at each position. To account for this, Margin implements a pruning and merging heuristic where the input is divided into smaller pieces, unlikely states are pruned, and the resulting graphs are merged before running the full forward-backward algorithm.

#### Improved Functionality

One of the most significant changes to the Margin workflow is that we now only analyze sites proposed by the input VCF. Previously we considered any reference position where less than 80% of the nucleotides agreed with the reference base as a candidate variant site. To determine which proposed variants are considered, we read the input VCF and remove all INDEL variants (`phase.onlyUseSNPVCFEntries = false`), all homozygous variants (`phase.includeHomozygousVCFEntries = false`), all non-PASS variants (`phase.onlyUsePassVCFEntries = true`), and all variants with a quality score below `phase.minVariantQuality = 10`. For each chunk, we perform an adaptive sampling of variants (`phase.useVariantSelectionAdaptiveSampling = true`) where we start by taking all variants with a quality score above `phase.variantSelectionAdaptiveSamplingPrimaryThreshold = 20`. If the average distance between these variants for the chunk is greater than `phase. variantSelectionAdaptiveSamplingDesiredBasepairsPerVariant = 2000`, we take variants ordered by quality score descending until we achieve the desired number of variants. These values were determined after experimentation on HG002.

Instead of considering only the nucleotide aligned directly to the variant position as the first iteration of Margin had done, we now extract substrings from the reference and reads and perform an alignment to determine which allele the read most likely originated from. In theory, this allows Margin to use INDELs during phasing, although for our current evaluations we do not test this functionality. When extracting reference bases, we take `phase.columnAnchorTrim = 12` bp from the reference before the variant position, and the same amount upstream from the end of the variant position. All alleles (including the reference allele) in the VCF are recreated by substituting the replaced reference base with the allelic sequence from the VCF. Note that there is no further modification of the reference sequences for cases where multiple variants fall within the extracted region. From the reads, we extract all sequence between the first and last position matched to the reference over the extracted reference region. Reads substrings for which the average base-level quality score over the substring is less than `polish.minAvgBaseQuality = 10` are excluded at these positions.

The original Margin phasing HMM used an emission model based off of a read error model (empirically determined) and a mutation likelihood model (based on likelihood of specific mutations). In the new implementation, we replace this with a likelihood generated by aligning the read substrings to the allelic substrings.

As in the previous iteration, after running the forward-backward algorithm on the HMM we marginalize over possible genotypes at each variant site to determine final predicted haplotypes. Given these haplotypes and the read-to-allele alignment likelihoods, we compute from which haplotype the read most likely originated using the joint probability of the read substrings aligning to the alleles for each haplotype. This haplotyping step is performed for all reads used for phasing, as well as other reads filtered out before phasing (as described below in the Chunking section). If a read does not span any variants or has equal likelihood of aligning to both haplotypes, no prediction is made regarding the haplotype from which it originated.

#### Chunking

Margin divides input into `polish.chunkSize = 100000` bp chunks with `polish.chunkBoundary = 10000` bp boundary on both ends, resulting in 2x `polish.chunkBoundary` overlap between chunks. Once the boundaries have been determined, the ordering of the chunks is mutated (`polish.shuffleChunks = true`) by default based on descending order of size (`polish.shuffleChunksMethod = size_desc`), with random ordering (`random`) also as a configurable option. While the ordering does not have a large effect on the runtime during phasing, we found that deep chunks would take drastically longer to complete for polishing, and ensuring that they were completed first would reduce overall runtime for the submodule.

When operating on a chunk, Margin first extracts all the reads that have an alignment between the start and end of the chunk, tracking all alignments falling between the start and end of the extended boundary. Margin then collects a set of reads to run the main algorithm on, first by removing reads that have a MQ score below `polish.filterAlignmentsWithMapQBelowThisThreshold = 10`, are secondary alignments (`polish.includeSecondaryAlignments = false`), or are supplementary alignments (`polish.includeSupplementaryAlignments = false`). Given this set of reads, we downsample to an expected depth of `polish.maxDepth = 64` for haplotagging and `polish.maxDepth = 32` for variant phasing. The downsampling is biased to maximize coverage over heterozygous variants given the constraint on expected coverage. To accomplish this, we compute the sampling probabilities for each read according to the following linear program, which we solve using the LP Solve library (http://lpsolve.sourceforge.net/).

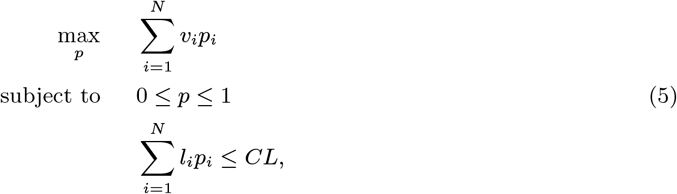

where *p_i_* is the probability of selecting read *i*, *v_i_* is the number of heterozygous variants on read *i*, *l_i_* is the length of read *i*, *C* is the desired coverage, and *L* is the length of the chunk.

The algorithm runs on the chunk region with the primary set of reads (all reads kept after filtering and downsampling) and assigns reads and variants to haplotypes. Then it takes any removed reads (either through filtering or downsampling) and assigns them to haplotypes as described above. Margin tracks the assignment of variants to haplotypes within the chunk (not including the boundaries), and of reads to haplotypes for the whole region (including the boundaries).

To stitch two chunks together, we need to determine whether the two previous haplotypes (*P*_1_,*P*_2_) are oriented with the two current haplotypes (*C*_1_, *C*_2_) in *cis* (*P*_1_*C*_1_, *P*_2_*C*_2_) or in *trans* (*P*_1_*C*_2_, *P*_2_*C*_1_). To do this, we compare the number of reads that are in *cis* and in *trans* between the two chunks; if there are more reads in *trans*, we switch the haplotypes of the current chunk’s reads and variants. To mulithread this process, Margin separates all the chunks in a contig into `numThreads` contiguous groups. The chunks in each of the groups are stitched together by a single thread, and then the same stitching process is used to stitch each of these groups together to complete the whole contig.

The final assignment of a read to a haplotype is determined by the haplotype it was assigned to in the first chunk for the contig. The final assignment of an allele to a haplotype is determined by the chunk it falls within (boundary region excluded).

#### Phaseset Determination

When writing the output VCF, Margin makes predictions about which sets of variants are confidently inherited together and annotates the output with phaseset (PS) tags. Margin will assign a phaseset to a variant if it is heterozygous, a SNP (`phase.onlyUseSNPVCFEntries = true`), and if it agrees with the genotype from the input VCF (`phase.updateAllOutputVCFFormatFields = false`). For variants meeting this criteria, as Margin iterates through the VCF it will extend the current phaseset unless *(a)* the variant is the first in the contig, *(b)* there are no reads spanning between the current variant and the previous variant, *(c)* there is an unlikely division of reads for the variant (explained below), or *(d)* the reads spanning the current variant and the previous variant are discordant above some threshold (explained below). The values described below were determined after experimentation on HG002.

To identify unlikely divisions of reads (which we take as potential evidence there is an error in the phasing), we take the number of primary reads assigned to each haplotype and find the binomial p-value for that division of reads. If that probability is less than the threshold `phase.phasesetMinBinomialReadSplitLikelihood = 0.0000001`, we create a new phase set for this variant.

Within each chunk region (boundary region excluded) and after determining haplotype assignment for the reads, we track which primary reads were used for phasing and to which haplotype they were assigned in the chunk. This serves as a check against poorly-phased or poorly-stitched chunks. To determine discordancy in the phasing between variants, we compare the number of reads which are in *cis* or “concordant” *(C_c_*) given the read assignment to adjacent variants, and the number in *trans* or “discordant” *(C_d_)* between variants. If the discordancy ratio *C_d_*/(*C_c_* + *C_d_*) is greater than `phase.phasesetMaxDiscordantRatio = 0.5`, we create a new phase set for this variant.

#### Margin: Code availability

Margin is available at https://github.com/UCSC-nanopore-cgl/margin.

### Local Phasing Correctness

The local phasing correctness (LPC) is a novel metric for measuring phasing accuracy that we developed for this study. More precisely, the LPC is a family of metrics parameterized by a varying parameter *ρ* ∈ [0,1], which controls the degree of locality. The LPC can be seen as a generalization of the two most common metrics used to evaluate phasing accuracy: the switch error rate and the Hamming rate. The switch error rate corresponds to the LPC with *ρ* = 0 (fully local), and the Hamming rate is closely related to the LPC with *ρ* = 1 (fully global). With intermediate values of *ρ*, the LPC can measure meso-level phasing accuracy that the two existing metrics cannot quantify.

The LPC consists of a sum over all pairs of heterozygous variants where each pair contributes an amount that decays with greater genomic distance. If the variants are incorrectly phased relative to each other, the pair contributes 0. This sum is normalized by its maximum value so that the LPC is always takes a value between 0 and 1. In mathematical notation,

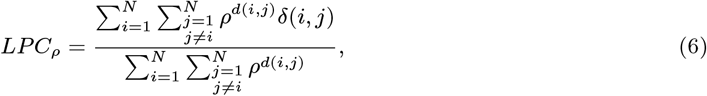

where *d*(*i,j*) is the distance between variants *i* and *j*, and *δ*(*i,j*) is an indicator for whether variants *i* and *j* are correctly phased relative to each other. In the case that *ρ* = 0 the above formula is undefined, and we instead take the limit of *LPC_ρ_* as *ρ* → 0.

If we take *d*(*i,j*) to be |*i* — *j*|, then it can be shown that *LPC*_0_ is equivalent to the complement of the switch error rate (i.e. the “switch correctness” rate). *LPC*_1_ is not equivalent to the Hamming rate, but they are monotonic functions of each other. Thus, *LPC*_1_ and Hamming rate always produce the same relative ranking but not the same numerical value. Alternatively, we can take *d*(*i,j*) to be the genomic distance between the variants, measured in base pairs. In doing so, the LPC no longer has the provable relationships to switch error rate and Hamming rate, neither of which has any mechanism to incorporate genomic distance. However, we consider the genomic distance to be a more relevant measurement of distance for phasing accuracy than the variants’ ordinal numbers. Accordingly, we believe that this amounts to a further strength of the LPC over existing metrics, and all LPC values reported in this work use this definition of distance.

The *ρ* parameter is mathematically convenient but difficult to interpret. To improve interpretability, we can reparameterize the LPC with *λ* = — log 2/ log *ρ*, which we call the “length scale”. This is the distance (measured either in base pairs or number of variants) at which a pair of variants has 1/2 of the maximum weight. The length scale gives an approximate sense of the scale of distances that the LPC incorporates, although it is worth stressing that pairs that are closer together always receive more weight than pairs that are further apart.

#### Local Phasing Correctness: Code availability

The code used to calculate LPC is available at https://github.com/UCSC-nanopore-cgl/margin.

### DeepVariant

DeepVariant is a small variant identification method based on deep neural networks that achieve high performance on different short-read and long-read sequencing platforms [36]. DeepVariant has released models for different datatypes and a details of the methods implemented in DeepVariant can be found in associated releases [36, 21]. Here we present an overview of the methods we implemented to achieve high performance with nanopore data.

### Adapting DeepVariant to Oxford Nanopore reads

DeepVariant performs variant calling in three stages, “make examples”, “call variants”, and “postprocess variants”. In “make examples”, potential variant positions are identified by applying a minimal threshold for evidence. In positions meeting the candidate generation criteria, the reads overlapping the position are converted into a pileup of a 221-bp window centered at the variant. Multiple features of the reads are represented as different dimensions in the pileup, including the read base, base quality, mapping quality, strand, whether the read supports the variant, and whether the base matches the reference. Prior to this work, the heuristics for candidate generation were simple (at least 2 reads supporting a variant allele and a variant allele fraction at least 0.12). However, the higher error rate of Oxford Nanopore data generated far too many candidates for DeepVariant to call variants in a genome in a reasonable time. To integrate DeepVariant with PEPPER, we developed the ability to import candidates directly from a VCF to replace the logic in “make examples”. This allows DeepVariant to read the output of PEPPER, and in theory, can be used in a similar manner with the outputs of other methods.

DeepVariant has released models for different datatypes [36, 21]. In order to achieve high performance on Oxford Nanopore data, a new DeepVariant model trained for this datatype was required. For this, we modified the training process for DeepVariant. The training process for DeepVariant is very similar to the variant calling process (generating candidates from make examples in a similar manner), but with the addition of a step which reconciles a candidate variant with the truth label from Genome-in-a-Bottle. We adapted this process to use the `VcfCandidateImporter` to propose PEPPER candidates. Because the representations of variants can diverge from the representation in GIAB (even for the same event), modifying the process for training required multiple rounds of iteration to identify mislabeled edge cases and to modify the proposed candidate representation for training.

With these modifications, training of DeepVariant models using the existing machinery could proceed. These modifications also allowed existing logic for the stages “call variants” and “postprocess variants” to work directly with the trained Oxford Nanopore model.

#### DeepVariant: Code availability

All code in the DeepVariant repository incorporates the improvements made for this paper (https://github.com/google/deepvariant). This repository also contains a model retraining tutorial (https://github.com/google/deepvariant/blob/r1.1/docs/deepvariant-training-case-study.md).

### Assembly polishing with PEPPER-Margin-DeepVariant

The assembly polishing method of PEPPER-Margin-DeepVariant is described below:

1. **PEPPER-SNP**: PEPPER-SNP finds single nucleotide polymorphisms (SNPs) from the read alignments to a haploid assembly using a RNN. For assembly polishing, we use the same infrastructure described in the haplotype-aware variant calling section.
2. **Margin**: Margin takes the SNPs reported by PEPPER-SNP and generates a haplotagged alignment file using a HMM. For assembly polishing, we use the same infrastructure described in the haplotype-aware variant calling section.
3. **PEPPER-HP**: PEPPER-HP takes the haplotagged alignment file and evaluates each haplotype independently and produces haplotype-specific candidate SNP and INDEL-like errors present in the assembly.

- *Haplotype-1 and Haplotype-2 candidate finding:* For haplotype-1, we take reads with `HP-1` and `HP-0` and generate base-level summary statistics. Then we use a RNN to produce nucleotide base predictions for each location of the genome. Finally, we find all the observed SNP or INDEL-like candidates in the reads with `HP-1` and `HP-0` tag and calculate a likelihood of each candidate to be a potential error in the assembly. If the likelihood of the candidate is above a set threshold, we propose that candidate as a potential edit for haplotype-1(`HP-1`) of the assembly. For haplotype-2, we take reads with `HP-2` and `HP-0` and find SNP and INDEL-like candidate errors following the same process described in Haplotype-1 candidate finding.
4. **DeepVariant**: DeepVariant takes the haplotype-specific candidate set from PEPPER-HP and the haplotagged alignment from Margin to identify errors present in the assembly using a convolutional neural network (CNN).

- *Haplotype-1 and haplotype-2 polishing:* For each candidate error from haplotype-1 set proposed by PEPPER-HP, we generate a feature set representing base, base-quality, mapping quality etc. in a tensor like format. In haplotype-specific polishing setup, reads are sorted by their haplotags in the order `{HP-1, HP-0, HP-2}`. Then we use a pre-trained `inception_v3` CNN to generate a prediction of `{0/0, 1/1}` for each candidate. Candidates in haplotype-1 that are classified as `{1/1}` are considered as missing heterozygous variants from the haploid assembly or an error present in the assembly. We apply the candidates of haplotype-1 classified as `{1/1}` to the haploid assembly using bcftools consensus to get a polished assembly `haplotype_1.fasta` representing one haplotype of the sample. For haplotype-2, we sort the reads by their haplotags in the order `{HP-2, HP-0, HP-1}`. Can-didates in haplotype-2 that are classified as `{1/1}` are similarly considered to be a missing heterozygous variant or an error present in the assembly, and are applied to the haploid assembly to get the second polished haplotype (`haplotype_2.fasta`).

We trained the PEPPER-Margin-DeepVariant assembly polishing method with HG002 assembly generated by the Shasta assembler. We first aligned the HG002 Shasta assembly to GRCh38 reference with dipcall[39] to associate assembly contigs with GRCh38 reference. Then we aligned HG002 GIAB v4.2.1 small variant benchmarking set to the assembly and marked the high-confidence regions to restrict training only in regions that GIAB notes as high-quality. The alignment between HG002 GIAB v4.2.1 to the assembly produced the training set. We trained PEPPER and DeepVariant on contigs that associate to `chr1-chr19` and used `chr20` as a holdout set. We used the same approach for Oxford Nanopore and PacBio HiFi based models.

