## Supplementary Results for "Haplotype-aware variant calling enables high accuracy in nanopore long-reads using deep neural networks"

#### Supplementary Notes

##### Data availability

###### Analysis data

We have made all of our analysis outputs i.e. variant calling VCF, phased VCF, and genome assemblies publicly available:
https://console.cloud.google.com/storage/browser/pepper-deepvariant-public/analysis_data/

###### Reference Sequences

We have used `GRCh38` and `GRCh37` human genome reference sequence that are publicly available:

`GRCh38`: ftp://ftp.ncbi.nlm.nih.gov/genomes/all/GCA/000/001/405/GCA_000001405.15_GRCh38/seqs_for_alignment_pipelines.ucsc_ids/GCA_000001405.15_GRCh38_no_alt_analysis_set.fna.gz

`GRCh37`: ftp://ftp-trace.ncbi.nih.gov/1000genomes/ftp/technical/reference/phase2_reference_assembly_sequence/hs37d5.fa.gz

###### Oxford Nanopore Sequencing data

We used publicly available data of ten samples to evaluate different methods. Following are the links to the fastq files that we used:

`HG001`: https://s3-us-west-2.amazonaws.com/human-pangenomics/index.html?prefix=NHGRI_UCSC_panel/HG001/nanopore/Guppy_4.2.2/

- `HG001_Circulomics_Guppy_4.2.2.fastq.gz`
- `HG001_NBT2018_Guppy_4.2.2.fastq.gz`

`HG002`: https://s3-us-west-2.amazonaws.com/human-pangenomics/index.html?prefix=NHGRI_UCSC_panel/HG002/nanopore/Guppy_4.2.2/

- `GM24385_1_Guppy_4.2.2_prom.fastq.gz`
- `GM24385_2_Guppy_4.2.2_prom.fastq.gz`
- `GM24385_3_Guppy_4.2.2_prom.fastq.gz`

`HG003`: https://s3-us-west-2.amazonaws.com/human-pangenomics/index.html?prefix=NHGRI_UCSC_panel/HG003/nanopore/Guppy_4.2.2/

- `GM24149_1_Guppy_4.2.2_prom.fastq.gz`
- `GM24149_2_Guppy_4.2.2_prom.fastq.gz`
- `GM24149_3_Guppy_4.2.2_prom.fastq.gz`

`HG004`: https://s3-us-west-2.amazonaws.com/human-pangenomics/index.html?prefix=NHGRI_UCSC_panel/HG004/nanopore/Guppy_4.2.2/

- GM24143_1_Guppy_4.2.2_prom.fastq.gz
- GM24143_2_Guppy_4.2.2_prom.fastq.gz
- GM24143_3_Guppy_4.2.2_prom.fastq.gz

HG005: https://s3-us-west-2.amazonaws.com/human-pangenomics/index.html?prefix=NHGRI_UCSC_panel/HG005/nanopore/Guppy_4.2.2/

- 01_09_20_R941_GM24631_1_Guppy_4.2.2_prom.fastq.gz
- 01_09_20_R941_GM24631_2_Guppy_4.2.2_prom.fastq.gz
- 01_09_20_R941_GM24631_3_Guppy_4.2.2_prom.fastq.gz

HG006: https://s3-us-west-2.amazonaws.com/human-pangenomics/index.html?prefix=NHGRI_UCSC_panel/HG006/nanopore/Guppy_4.2.2/

- 01_09_20_R941_GM24694_1_Guppy_4.2.2_prom.fastq.gz
- 01_09_20_R941_GM24694_2_Guppy_4.2.2_prom.fastq.gz
- 01_09_20_R941_GM24694_3_Guppy_4.2.2_prom.fastq.gz

HG007: https://s3-us-west-2.amazonaws.com/human-pangenomics/index.html?prefix=NHGRI_UCSC_panel/HG007/nanopore/Guppy_4.2.2/

- 01_09_20_R941_GM24695_1_Guppy_4.2.2_prom.fastq.gz
- 01_09_20_R941_GM24695_2_Guppy_4.2.2_prom.fastq.gz
- 01_09_20_R941_GM24695_3_Guppy_4.2.2_prom.fastq.gz

HG00733: https://s3-us-west-2.amazonaws.com/human-pangenomics/index.html?prefix=NHGRI_UCSC_panel/HG00733/nanopore/Guppy_4.2.2/

- HG00733_1_Guppy_4.2.2_prom.fastq.gz
- HG00733_2_Guppy_4.2.2_prom.fastq.gz
- HG00733_3_Guppy_4.2.2_prom.fastq.gz

HG02723: https://s3-us-west-2.amazonaws.com/human-pangenomics/index.html?prefix=NHGRI_UCSC_panel/HG02723/nanopore/Guppy_4.2.2/

- HG02723_1_Guppy_4.2.2_prom.fastq.gz
- HG02723_2_Guppy_4.2.2_prom.fastq.gz
- HG02723_2_Guppy_4.2.2_prom.fastq.gz

CHM13: https://s3.amazonaws.com/nanopore-human-wgs/chm13/nanopore/rel6/rel6.fastq.gz

##### PacBio HiFi sequencing data

We used publicly available PacBio HiFi sequencing data of eight samples to evaluate different methods. Following are the links to the data locations:

HG001: ftp://ftp-trace.ncbi.nlm.nih.gov/giab/ftp/data/NA12878/

HG002: https://storage.googleapis.com/pepper-deepvariant-public/sequencing_data/PacBio_HiFi/HG002/HG002_35x_PacBio_14kb-15kb.fastq.gz

HG003: https://storage.googleapis.com/pepper-deepvariant-public/sequencing_data/PacBio_HiFi/HG003/HG003_35x_PacBio_14kb-15kb.fastq.gz

HG004: https://storage.googleapis.com/pepper-deepvariant-public/sequencing_data/PacBio_HiFi/HG004/HG004_35x_PacBio_14kb-15kb.fastq.gz

HG005: https://s3-us-west-2.amazonaws.com/human-pangenomics/index.html?prefix=submissions/8ced218e-9699-458e-9708-eef6969a8065--EXTRAMURAL_SAMPLES/HG005/PacBio_HiFi/

- PBmixSequel788_1_A01_PBXX_30hours_19kbV2PD_70pM_HumanHG005_CCS/m64109_200304_195708.fastq.gz

- PBmixSequel789_1_A01_PBXW_30hours_15kbV2PD_70pM_HumanHG005_CCS/m64109_200309_192110.fastq.gz
- PBmixSequel789_2_B01_PBXX_30hours_19kbV2PD_70pM_HumanHG005_CCS/m64109_200311_013444.fastq.gz
- PBmixSequel840_1_A01_PCCD_30hours_15kbV2PD_70pM_HumanHG005_CCS/m64017_200723_190224.fastq.gz
- PBmixSequel842_1_A01_PCCD_30hours_15kbV2PD_70pM_HumanHG005_CCS/m64017_200730_190124.fastq.gz
- PBmixSequel842_2_B01_PCCD_30hours_15kbV2PD_70pM_HumanHG005_CCS/m64017_200801_011415.fastq.gz
- PBmixSequel842_3_C01_PCCD_30hours_15kbV2PD_70pM_HumanHG005_CCS/m64017_200802_073944.fastq.gz

HG00733: https://s3-us-west-2.amazonaws.com/human-pangenomics/index.html?prefix=submissions/8ced218e-9699-458e-9708-eef6969a8065--EXTRAMURAL_SAMPLES/HG00733/hifi/

- HG00733_20190925_EEE_m54329U_190607_185248.Q20.fastq.gz
- HG00733_20190925_EEE_m54329U_190615_010947.Q20.fastq.gz
- HG00733_20190925_EEE_m54329U_190617_231905.Q20.fastq.gz
- HG00733_20190925_EEE_m54329U_190619_052546.Q20.fastq.gz
- HG00733_20190925_EEE_m54329U_190629_180018.Q20.fastq.gz
- HG00733_20190925_EEE_m54329U_190701_222759.Q20.fastq.gz
- HG00733_20190925_EEE_m54329U_190827_173812.Q20.fastq.gz

HG02723: https://s3-us-west-2.amazonaws.com/human-pangenomics/index.html?prefix=submissions/8fa7bde9-be6f-4160-97a9-b639a8962c66--WUSTL_OTHER_HiFi/HG02723/PacBio_HiFi/

- m64043_191221_024136.ccs.bam
- m64043_191223_180311.ccs.bam
- m64043_191225_001554.ccs.bam
- m64043_191226_064057.ccs.bam

CHM13: https://github.com/nanopore-wgs-consortium/CHM13/

**Illumina sequencing data**

We used publicly available Illumina short read data of three samples to perform variant calling assessment.
HG002:

- https://storage.googleapis.com/pepper-deepvariant-public/sequencing_data/Illumina_novaseq/HG002/HG002.novaseq.pcr-free.35x.R1.fastq.gz
- https://storage.googleapis.com/pepper-deepvariant-public/sequencing_data/Illumina_novaseq/HG002/HG002.novaseq.pcr-free.35x.R2.fastq.gz

HG003:

- https://storage.googleapis.com/pepper-deepvariant-public/sequencing_data/Illumina_novaseq/HG003/HG003.novaseq.pcr-free.35x.R1.fastq.gz
- https://storage.googleapis.com/pepper-deepvariant-public/sequencing_data/Illumina_novaseq/HG003/HG003.novaseq.pcr-free.35x.R2.fastq.gz

HG004:

- https://storage.googleapis.com/pepper-deepvariant-public/sequencing_data/Illumina_novaseq/HG004/HG004.novaseq.pcr-free.35x.R1.fastq.gz
- https://storage.googleapis.com/pepper-deepvariant-public/sequencing_data/Illumina_novaseq/HG004/HG004.novaseq.pcr-free.35x.R2.fastq.gz

We used publicly available Illumina short read data for assembly quality assessment.
`HG005-HG006-HG007:`

- https://s3-us-west-2.amazonaws.com/human-pangenomics/index.html?prefix=submissions/00c8916e-a869-473a-9a03-e39867b35499--HG005_CHILD/HG005/Illumina/
- https://s3-us-west-2.amazonaws.com/human-pangenomics/index.html?prefix=submissions/8ced218e-9699-458e-9708-eef6969a8065--EXTRAMURAL_SAMPLES/HG005/parents/HG006/Illumina/
- https://s3-us-west-2.amazonaws.com/human-pangenomics/index.html?prefix=submissions/8ced218e-9699-458e-9708-eef6969a8065--EXTRAMURAL_SAMPLES/HG005/parents/HG007/Illumina/

`HG00731-HG00732-HG00733:`

- https://s3-us-west-2.amazonaws.com/human-pangenomics/index.html?prefix=submissions/325b4b1c-9f20-49be-b03a-596da89c466e--1000G_CHILDREN/HG00733/1000G_data/HG00733.final.cram
- https://s3-us-west-2.amazonaws.com/human-pangenomics/index.html?prefix=submissions/54D542ED-6650-4A12-8B40-0F92CC320486--1000G_parents/HG00733/parents/HG00732/HG00732.final.cram
- https://s3-us-west-2.amazonaws.com/human-pangenomics/index.html?prefix=submissions/54D542ED-6650-4A12-8B40-0F92CC320486--1000G_parents/HG00733/parents/HG00731/HG00731.final.cram

`HG02721-HG02722-HG02723:`

- https://s3-us-west-2.amazonaws.com/human-pangenomics/index.html?prefix=submissions/325b4b1c-9f20-49be-b03a-596da89c466e--1000G_CHILDREN/HG02723/1000G_data/HG02723.final.cram
- https://s3-us-west-2.amazonaws.com/human-pangenomics/index.html?prefix=submissions/54D542ED-6650-4A12-8B40-0F92CC320486--1000G_parents/HG02723/parents/HG02722/HG02722.final.cram
- https://s3-us-west-2.amazonaws.com/human-pangenomics/index.html?prefix=submissions/54D542ED-6650-4A12-8B40-0F92CC320486--1000G_parents/HG02723/parents/HG02721/HG02721.final.cram

`CHM13:` https://github.com/nanopore-wgs-consortium/CHM13

##### Genome-In-A-Bottle (GIAB) benchmakring data

We used Genome-In-A-Bottle (GIAB) truth set for seven genomes HG001-HG007. For each genome, we have a VCF file describing the truth variants and a bed file describing regions to use for benchmarking. Both these files are input to `hap.py` when benchmarking a query VCF. In this truth set, `v3.3.2` was generated for `GRCh37` reference and `v4.2.1` was generated for `GRCh38` human reference genome. While evaluating our pipeline, we used associated reference while generating the benchmarking results.

The GIAB benchmarking data is publicly available:
`HG001`
ftp://ftp-trace.ncbi.nlm.nih.gov/giab/ftp/release/NA12878_HG001/NISTv3.3.2/GRCh37/
`HG002`
ftp://ftp-trace.ncbi.nlm.nih.gov/giab/ftp/release/AshkenazimTrio/HG002_NA24385_son/NISTv4.2.1/GRCh38/
`HG003`
ftp://ftp-trace.ncbi.nlm.nih.gov/giab/ftp/release/AshkenazimTrio/HG003_NA24149_father/NISTv4.2.1/GRCh38/
`HG004`
ftp://ftp-trace.ncbi.nlm.nih.gov/giab/ftp/release/AshkenazimTrio/HG004_NA24143_mother/NISTv4.2.1/GRCh38/
`HG005`
ftp://ftp-trace.ncbi.nlm.nih.gov/giab/ftp/release/ChineseTrio/HG005_NA24631_son/NISTv3.3.2/GRCh37/

`HG006`
ftp://ftp-trace.ncbi.nlm.nih.gov/giab/ftp/release/ChineseTrio/HG006_NA24694_father/NISTv3.3.2/GRCh37/
`HG007`
ftp://ftp-trace.ncbi.nlm.nih.gov/giab/ftp/release/ChineseTrio/HG007_NA24695_mother/NISTv3.3.2/GRCh37/
`Stratification regions of GRCh38`
https://ftp-trace.ncbi.nlm.nih.gov/giab/ftp/release/genome-stratifications/v2.0/

##### Gencode annotation files

We used gencode version `v35` annotation of `GRCh38` and `GRCh37` to determine variant calling accuracy in gene regions. The gencode annotation file is publicly available:

`Gencode v35`
ftp://ftp.ebi.ac.uk/pub/databases/gencode/Gencode_human/release_35/gencode.v35.annotation.gtf.gz

##### Execution Parameters

##### Minimap2, PBMM2

`Minimap2` version `2.17-r941` from `https://github.com/lh3/minimap2`. We used `samtools` version `1.10` for sorting and filtering.

We used the following command to map nanopore reads with minimap2:

```
minimap2 -ax map-ont -t 32 \
  Reference.fasta \
  reads.fastq.gz | samtools view -hb -F 0x904 > unsorted.bam
samtools sort -@32 -o sorted.bam unsorted.bam
samtools index -@32 sorted.bam
```

For PacBio HiFi data we used `pbmm2` version `1.4.0` form `https://github.com/PacificBiosciences/pbmm2`. We used the following command to map PacBio HiFi reads with pbmm2:

```
docker run --ipc=host \
  -v /data:/data \
  quay.io/biocontainers/pbmm2:1.4.0--h56fc30b_0 pbmm2 align \
  --sort -j <THREADS> -J <THREADS> \
  --preset CCS \
  /data/reference.fa /data/reads.fastq /data/reads_2_ref.bam
```

##### Subsampling with samtools

We used `samtools` version `1.10` to subsample the alignment files at different coverages. We used the following command:

```
    samtools view -s 0.FRAC -@<THREDS> -b INPUT.bam > OUTPUT_DOWNSAMPLED.sam
```

##### PEPPER-Margin-DeepVariant variant calling

We used the following command to run the PEPPER-Margin-DeepVariant variant calling pipeline:

```
docker run --ipc=host \
  -v /data:/data \
  -u (id -u $USER):(id -g $USER) \
  kishwars/pepper_deepvariant:r0.4 \
  run_pepper_margin_deepvariant call_variant \
  -b <BAM> \
  -f <FASTA>\
  -o <OUTPUT_DIR> \
  -t <THREADS> \
  --ont # --ccs if using PacBio HiFi reads
```

When running the PEPPER-Margin-DeepVariant assembly polishing pipeline, we ran it with these arguments:

```
time docker run --ipc=host \
  -v /data:/data \
  kishwars/pepper_deepvariant:r0.4 \
```

```
  run_pepper_margin_deepvariant polish_assembly \
  -b <READS_2_ASSEMBLY_BAM> \
  -f <ASSEMBLY> \
  -o <OTUPUT_DIR> \
  -t <THREADS> \
  -s <SAMPLE_NAME>
  --ont # --ccs if using PacBio HiFi reads
```

**Medaka**

We used Medaka `v1.2.0` available from `https://github.com/nanoporetech/medaka`. We used the following command to variant call using Medaka:

```
medaka_variant -i INPUT.bam -f reference.fna -t THREADS -o OUTPUT_DIR/
```

**Clair**

We used Clair `v2.1.1` available from `https://github.com/HKU-BAL/Clair`. We used the following command to run `Clair`:

```
conda activate clair-env

python clair.py callVarBamParallel \
  --chkpnt_fn clair_model/ont/model \
  --ref_fn reference.fna \
  --bam_fn INPUT.bam \
  --threshold 0.2 \
  --sampleName SAMPLE \
  --output_prefix OUTPUT_DIR/OUTPUT > command_clair.sh

cat command_clair.sh | parallel -j16

for i in OUTPUT.*.vcf; do if ! [ -z "$(tail -c 1 "$i")" ]; then echo "$i"; fi ; done | \
grep -f - command.sh | sh

vcfcat OUTPUT.*.vcf | bcftools sort -m 2G | bgzip > OUTPUT.wgs.vcf.gz
bcftools filter -s LowQual -e '%QUAL<748' -Oz OUTPUT.wgs.vcf.gz > OUTPUT.wgs.FILT.vcf.gz
```

We evaluated the filtered file. The filter cutoff value is adopted from the github suggestion.

**Longshot**

We used longshot `v0.4.2` available from `https://github.com/pjedge/longshot`. We used the following command to run longshot:

```
longshot --strand_bias_pvalue_cutoff 0.01 \
  --bam INPUT.bam --ref reference.fna --out OUTPUT.VCF
```

**DeepVariant (Illumina and PacBio HiFi)**

We used `DeepVariant v1.1.0` available in `https://github.com/google/deepvariant` to produce variant calls using Illumina and PacBio HiFi data. Following are the commands we used. For PacBio HiFi variant calling, we used:

- **DeepVariant for Illumina**:

```
docker run \
  -v /data:/data \
  google/deepvariant:1.1.0 \
  /opt/deepvariant/bin/run_deepvariant \
  --model_type=WGS \
  --ref=/data/reference.fna \
  --reads=/data/INPUT.bam \
  --output_vcf=/data/OUTPUT.vcf.gz \
  --num_shards=THREADS
```

- **DeepVariant for PacBio HiFi**:

– Initial calling on unphased bam:

```
docker run \
  -v /data:/data \
  google/deepvariant:1.1.0 \
  /opt/deepvariant/bin/run_deepvariant \
  --model_type=PACBIO \
  --ref=/data/reference.fna \
  --reads=/data/INPUT.PHASED.bam \
  --output_vcf=/data/OUTPUT.vcf.gz \
  --num_shards=THREADS
```

– Haplotype-aware calling on phased bam:

```
docker run \
  -v /data:/data \
  google/deepvariant:1.1.0 \
  /opt/deepvariant/bin/run_deepvariant \
  --model_type=PACBIO \
  --ref=/data/reference.fna \
  --reads=/data/INPUT.PHASED.bam \
  --output_vcf=/data/OUTPUT.vcf.gz \
  --num_shards=THREADS \
  --use_hp_information
```

**Hap.py**

We used `hap.py` version `v0.3.12` to assess the variant calls against GIAB truth set. The `hap.py` program is available via `jmcdani20/hap.py:v0.3.12` docker image.

```
docker run -it -v /data/:/data/ \
  jmcdani20/hap.py:v0.3.12 /opt/hap.py/bin/hap.py \
  /data/GIAB_benchmark.vcf.gz \
  /data/INPUT.vcf.gz \
  -f /data/GIAB_benchmark.bed \
  -r /data/reference.fna \
  -o /data/output/prefix \
  --stratification /data/genome_stratification/v2.0-GRCh38-stratifications.tsv \
  --pass-only \
  --engine=vcfeval \
  --threads=<THREADS>
```

The parameters we used for `hap.py`:

- `GIAB_benchmark.vcf.gz`: GIAB truth VCF file.
- `INPUT.vcf.gz`: The input VCF to compare against GIAB.
- `-f`: GIAB regions defined for variant assessment.
- `-r`: Reference FASTA file (GRCh38 or GRCh37).
- `--stratification`: Stratification file list for generating stratified results.
- `--pass-only`: Only use the PASS variants from input for assessment.
- `--engine`: Comparison engine.
- `--threads`: Number of threads to use.

**Margin**

We used `Margin v2.0` available from `https://github.com/UCSC-nanopore-cgl/margin`.

To haplotag a bam, we used the following command:

```
margin phase \
  INPUT.bam \
  REFERENCE.fasta \
  INPUT.vcf \
  params/misc/allParams.XXX_haplotag.json \
  --threads THREAD_COUNT \
  --skipPhasedVCF
```

To phase a vcf, we used the following command:

```
margin phase \
  INPUT.bam \
  REFERENCE.fasta \
  INPUT.vcf \
  params/misc/allParams.phase_vcf.json \
  --threads THREAD_COUNT \
  --skipHaplotypeBAM
```

##### WhatsHap

We used `WhatsHap v1.0` available from `https://github.com/whatshap/whatshap`. To phased a VCF file, we used the following command:

```
whatshap phase \
  -o PHASED.OUTPUT.vcf \
  --ignore-read-groups \
  --reference reference.fna \
  INPUT.vcf \
  INPUT.bam
```

To produce haplotags for reads, we used the following command:

```
whatshap haplotag \
  --reference $FASTA_FILE \
  --regions $contig \
  --ignore-read-groups \
  --output $PHASED_BAM_OUTPUT_DIR/$contig.phased.bam \
  $WHATSHAP_OUTPUT_DIR/$contig.phased.vcf.gz \
  $BAM_FILE
```

##### Local phasing correctness

We used the `calcLocalPhasingCorrectness` executable in `Margin v2.0` to calculate local phasing correctness. We used the following command to generate LPC statistics:

```
calcLocalPhasingCorrectness -m 1 -M 100000000 -d -s TRUTH.vcf.gz QUERY.vcf.gz > OUTPUT.tsv
```

We plotted the results of LPC with `plot_haplotagging_lpc.py` script found in the `https://github.com/tpesout/genomics_scripts` repository. We used the following commands to generate the plots:

```
plot_haplotagging_lpc.py -f OUTPUT_1.png -b \
  -i DATA-1_TOOL-1.tsv -i DATA-1_TOOL-2.tsv -i DATA-2_TOOL-1.tsv ..
plot_haplotagging_lpc.py -f OUTPUT_2.png -b -l \
  -i DATA-1_TOOL-1.tsv -i DATA-1_TOOL-2.tsv -i DATA-2_TOOL-1.tsv ..
plot_haplotagging_lpc.py -f OUTPUT_3.png -b -l 100000 -L 10  \
  -i DATA-1_TOOL-1.tsv -i DATA-1_TOOL-2.tsv -i DATA-2_TOOL-1.tsv ..
```

##### Trio-Binning

We used `canu` to trio-bin the reads to generate the admixture sample used in Fig 3.

The command used for Nanopore data was:

```
canu -haplotype -p trio_Binned -d trio_Binned_dir genomeSize=3.3g \
-haplotypeFather Father.fastq -haplotypeMother Mother.fastq \
-nanopore-raw nanopore.fastq
```

The command used for PacBio HiFi data was:

```
canu -haplotype -p trio_Binned -d trio_Binned_dir genomeSize=3.3g \
-haplotypeFather Father.fastq -haplotypeMother Mother.fastq -pacbio-hifi HiFi.fastq
```

**Natural switch error**

We calculated the natural switch was determined using `haplotagging_stats.py` available in `https://github.com/tpesout/genomics_scripts`. We used the following command to calculate the values:

```
haplotagging_stats.py \
  --input HAPLOTAGGED.bam \
  --truth_hap1 HAP1.read_ids.txt \
  --truth_hap2 HAP2.read_ids.txt \
  --threads THREAD_COUNT
```

**Natural switch visualization**

We used `compare_read_phasing_hapBam.py` available in `https://github.com/tpesout/genomics_scripts`. We used the following command to visualize the natural switch:

```
compare_read_phasing_hapBam.py  \
  -1 HAP1.read_idx.txt  \
  -2 HAP2.read_idx.txt  \
  -i INPUT.chr1.bam  \
  -p INPUT.phaseset.bed  \
  -F  \
  -d 100  \
  -N
```

**Shasta**

We used `Shasta` version `0.7.0` (commit `06a639d36d26a4203c0b934d6e63c719750c5398`) available from `https://github.com/chanzuckerberg/shasta` to generate nanopore-based *de novo* assemblies. We used the following command to run Shasta:

```
shasta --input /reads/SAMPLE_READS*.fasta  \
--conf shasta/conf/Nanopore-Sep2020.conf \
--memoryMode filesystem --memoryBacking 2M --assemblyDirectory SAMPLE_OUTPUT_DIR
```

**Flye**

We used `Flye` version `2.8.2` available from `https://github.com/fenderglass/Flye` to generate nanopore-based *de novo* assemblies. We invoked Flye the following way:

```
flye --nano-raw \
  file1.fastq.gz \
  file2.fastq.gz \
  file3.fastq.gz
  --out-dir FLYE_OUTPUT/
```

**Trio-Hifiasm**

We used `hifiasm` version `0.14` available from `https://github.com/chhylp123/hifiasm`. We also used `YAK` in this pipeline which is available in `https://github.com/lh3/yak`. We generated the assembly in two-steps:

**Step 1:** Generate maternal and paternal k-mer counts from paired-end short-reads from the parent samples:

```
yak count \
  -b37 -t16 -o pat.yak <(cat pat_1.fq.gz pat_2.fq.gz) < (cat pat_1.fq.gz pat_2.fq.gz)

yak count \
  -b37 -t16 -o mat.yak <(cat mat_1.fq.gz mat_2.fq.gz) < (cat mat_1.fq.gz mat_2.fq.gz)
```

**Step 2:** Generate the assembly using sample-specific PacBio HiFi reads (`SAMPLE-HiFi_reads.fa.gz`):

```
hifiasm -o SAMPLE.asm -t<THREADS> -1 pat.yak -2 mat.yak SAMPLE-HiFi_reads.fa.gz
```

A WDL workflow of this pipeline is available in
`https://github.com/human-pangenomics/hpp_production_workflows/`.

##### Homopolymer run-length analysis

We used `runLengthMatrix` module of `margin` to perform the homopolymer run-length analysis. We used the following command to generate the run-length statistics:

```
./runLengthMatrix \
  <READS_2_ASSEMBLY.BAM> \
  <ASSEMBLY.FASTA> \
  <margin/params/base_params.json> \
  -o <OUTPUT_PATH> \
  -t <THREADS> \
  -l <MAX_RUN_LENGTH>
```

##### Dipcall

We used `dipcall` available in https://github.com/lh3/dipcall to generate small variant call from the assemblies.

```
./run-dipcall \
OUTPUT_DIR/OUTPUT_PREFIX \
REFERENCE.fa \
Assembly.hp1.fa \
Assembly.hp2.fa \
-t THREADS > ASSEMBLY_2_REFERENCE.mak

make -j2 -f ASSEMBLY_2_REFERENCE.mak
```

#### Supplementary Figures

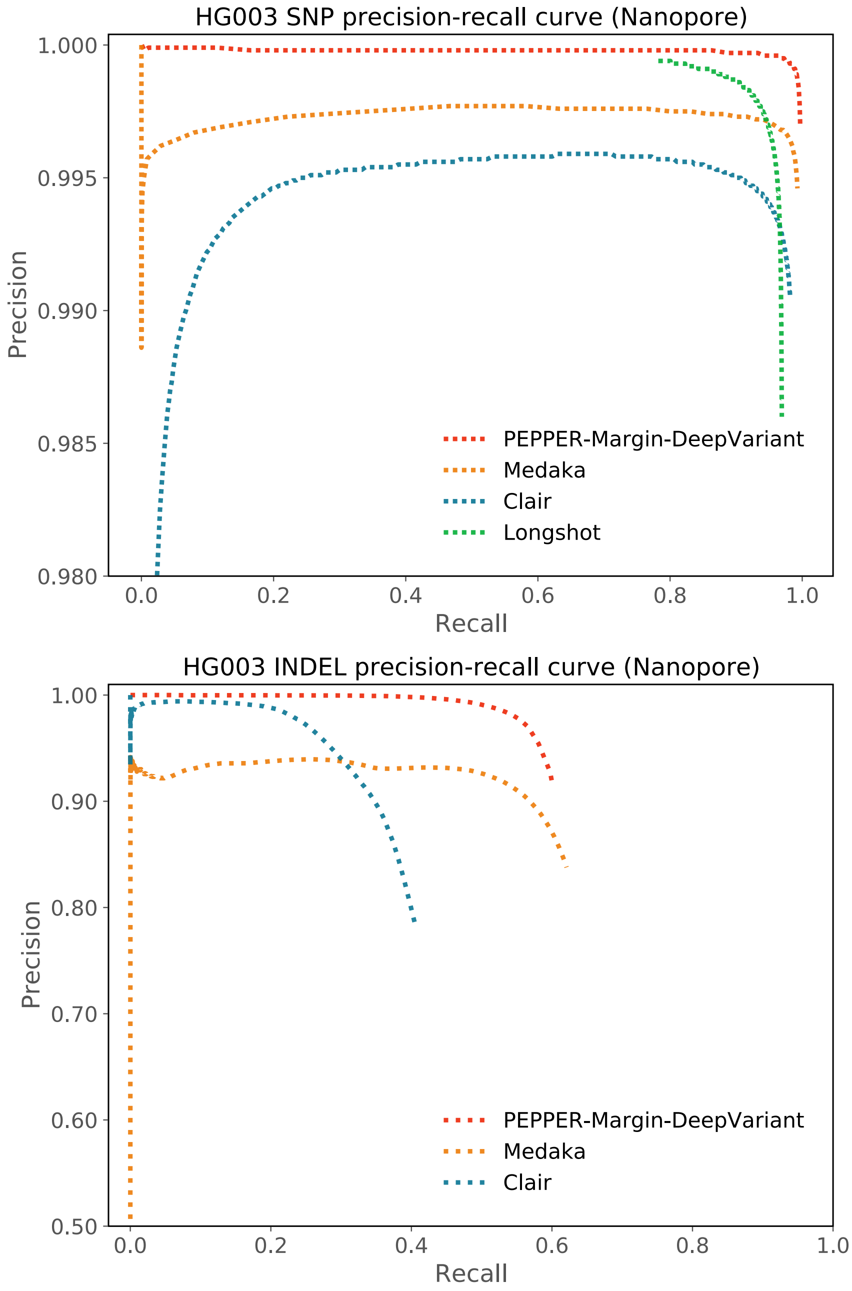


Supplementary Figure 1: Precision-Recall plot of HG003 for nanopore-based variant callers.

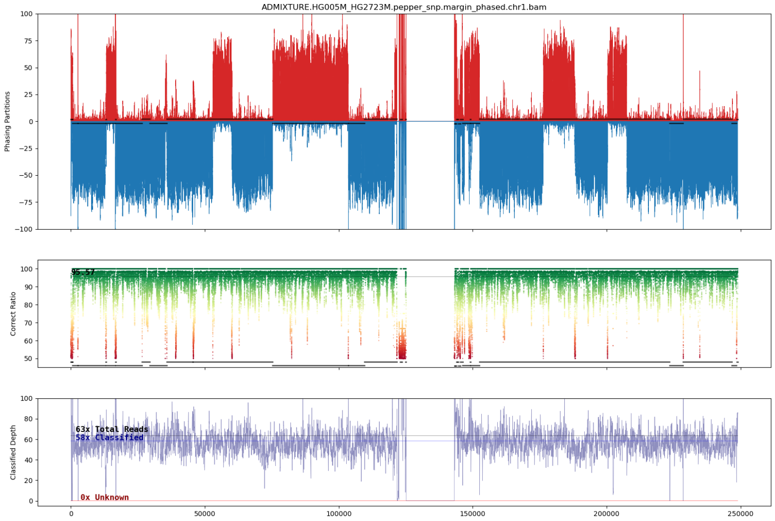


Supplementary Figure 2: Full Natural Switch plot for chr1 of an admixture of HG005 and HG02723's maternal haplotypes from nanopore data phased by Margin

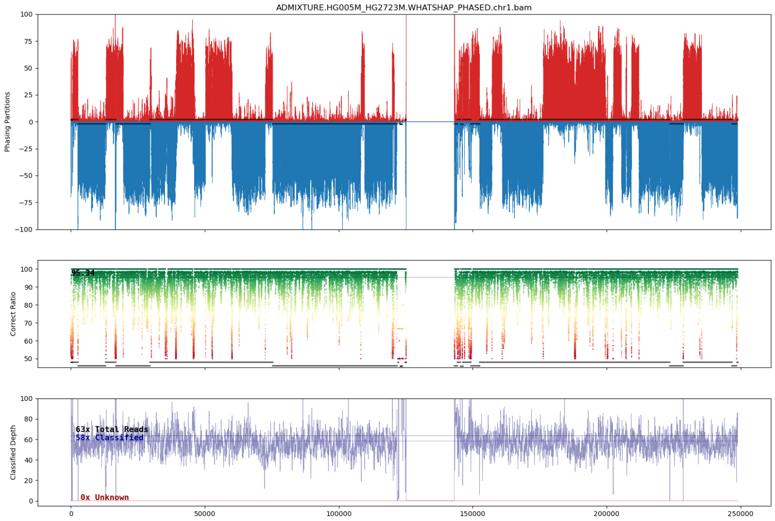


Supplementary Figure 3: Full Natural Switch plot for chr1 of an admixture of HG005 and HG02723's maternal haplotypes from nanopore data phased by WhatsHap

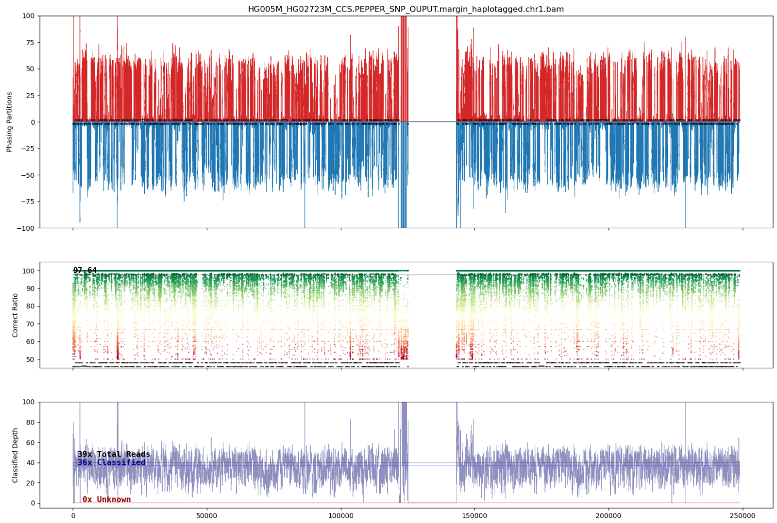


Supplementary Figure 4: Full Natural Switch plot for chr1 of an admixture of HG005 and HG02723's maternal haplotypes from PacBio HiFi data phased by Margin

### Supplementary Results

#### Variant calling results

| Sample Name | Type | Method | True positives | False negatives | False positives | Recall | Precision | F1-score |
|---|---|---|---|---|---|---|---|---|
| HG003 | SNP | P-M-DV | 3317032 | 10463 | 9958 | 0.9969 | 0.9970 | 0.9969 |
| | | Medaka | 3293174 | 22716 | 26549 | 0.9931 | 0.9920 | 0.9926 |
| | | Clair | 3266489 | 61006 | 31220 | 0.9817 | 0.9905 | 0.9861 |
| | | Longshot | 3224643 | 102852 | 45780 | 0.9691 | 0.9860 | 0.9775 |
| | INDEL | P-M-DV | 303643 | 200858 | 29400 | 0.6019 | 0.9136 | 0.7257 |
| | | Medaka | 313033 | 189639 | 69434 | 0.6227 | 0.8226 | 0.7089 |
| | | Clair | 205364 | 299137 | 58367 | 0.4071 | 0.7812 | 0.5352 |
| HG004 | SNP | P-M-DV | 3338882 | 7728 | 7474 | 0.9977 | 0.9978 | 0.9977 |
| | | Medaka | 3286457 | 20743 | 23309 | 0.9937 | 0.9930 | 0.9933 |
| | | Clair | 3285625 | 60985 | 32021 | 0.9818 | 0.9903 | 0.9860 |
| | | Longshot | 3243183 | 103427 | 45387 | 0.9691 | 0.9862 | 0.9776 |
| | INDEL | P-M-DV | 300258 | 210261 | 32429 | 0.5881 | 0.9046 | 0.7128 |
| | | Medaka | 306050 | 198390 | 88346 | 0.6067 | 0.7807 | 0.6828 |
| | | Clair | 203825 | 306694 | 61454 | 0.3993 | 0.7708 | 0.5260 |

Supplementary Table 1: Oxford nanopore variant calling performance comparison between Medaka, Clair, Longshot and PEPPER-Margin-DeepVariant (P-M-DV) on HG003 and HG004 with 90× coverage.

| HG003 coverage | Method | True positives | False negatives | False positives | Recall | Precision | F1-score (INDEL) |
|---|---|---|---|---|---|---|---|
| 20x | P-M-DV | 239619 | 264882 | 96644 | 0.4750 | 0.7164 | 0.5712 |
| | Medaka | 229787 | 268549 | 182403 | 0.4611 | 0.5628 | 0.5069 |
| | Clair | 142647 | 361854 | 40955 | 0.2827 | 0.7785 | 0.4148 |
| 30x | P-M-DV | 265318 | 239183 | 64214 | 0.5259 | 0.8083 | 0.6372 |
| | Medaka | 264877 | 233132 | 119190 | 0.5319 | 0.6949 | 0.6025 |
| | Clair | 167998 | 336503 | 44982 | 0.3330 | 0.7905 | 0.4686 |
| 40x | P-M-DV | 278902 | 225599 | 51381 | 0.5528 | 0.8472 | 0.6691 |
| | Medaka | 284431 | 217108 | 103944 | 0.5671 | 0.7373 | 0.6411 |
| | Clair | 180964 | 323537 | 48517 | 0.3587 | 0.7905 | 0.4935 |
| 50x | P-M-DV | 288480 | 216021 | 43169 | 0.5718 | 0.8723 | 0.6908 |
| | Medaka | 297390 | 206752 | 91135 | 0.5899 | 0.7702 | 0.6681 |
| | Clair | 189538 | 314963 | 51278 | 0.3757 | 0.7891 | 0.5090 |
| 60x | P-M-DV | 294414 | 210087 | 38056 | 0.5836 | 0.8878 | 0.7042 |
| | Medaka | 301161 | 198584 | 82479 | 0.6026 | 0.7896 | 0.6835 |
| | Clair | 195079 | 309422 | 53275 | 0.3867 | 0.7877 | 0.5187 |
| 70x | P-M-DV | 298553 | 205948 | 34079 | 0.5918 | 0.8997 | 0.7139 |
| | Medaka | 306842 | 194792 | 76519 | 0.6117 | 0.8048 | 0.6951 |
| | Clair | 199070 | 305431 | 55055 | 0.3946 | 0.7856 | 0.5253 |
| 80x | P-M-DV | 301312 | 203189 | 31269 | 0.5972 | 0.9079 | 0.7205 |
| | Medaka | 309376 | 191507 | 72591 | 0.6177 | 0.8142 | 0.7024 |
| | Clair | 202551 | 301950 | 56606 | 0.4015 | 0.7840 | 0.5310 |
| 90x | P-M-DV | 303643 | 200858 | 29400 | 0.6019 | 0.9136 | 0.7257 |
| | Medaka | 313033 | 189639 | 69434 | 0.6227 | 0.8226 | 0.7089 |
| | Clair | 205364 | 299137 | 58367 | 0.4071 | 0.7812 | 0.5352 |

Supplementary Table 2: Comparison on INDEL performance between Medaka, Clair and PEPPER-Margin-DeepVariant (P-M-DV) variant callers at different coverages of HG003 sample.

| Sample | Method | CPUs | Memory | GPUs | CPU cost/h | GPU cost/h | Instance cost/h | Total runtime | Total cost |
|---|---|---|---|---|---|---|---|---|---|
| HG001 50x ONT | Longshot | 2vCPUs | 13 GB | - | $0.12 | $0.00 | $0.12 | Out of memory | |
| | | 8vCPUs | 52 GB | - | $0.47 | $0.00 | $0.47 | Out of memory | |
| | | 16vCPUs | 104 GB | - | $0.95 | $0.00 | $0.95 | 51:25:31 | $48.84 |
| | Clair | 96vCPUs | 360 GB | - | $4.56 | $0.00 | $4.56 | 02:30:05 | $11.40 |
| | Medaka | 8vCPUs | 30GB | 1x NVIDIA Tesla P100 | $0.47 | $1.46 | $1.93 | Out of memory | |
| | | 16vCPUs | 104 GB | 1x NVIDIA Tesla P100 | $0.95 | $1.46 | $2.41 | 40:21:11 | $97.24 |
| | | 16vCPUs | 104 GB | - | $0.95 | $0.00 | $0.95 | 95:14:01 | $90.47 |
| | PEPPER Margin DeepVariant | 96vCPUs | 360 GB | - | $4.56 | $0.00 | $4.56 | 12:59:19 | $59.28 |
| | | 96vCPUs | 360 GB | 4x NVIDIA Tesla P100 | $4.56 | $5.84 | $10.4 | 6:41:56 | $70 |
| | | 2vCPUs | 13 GB | - | $0.12 | $0.00 | $0.12 | Out of memory | |

| HG003 coverage | Method | True positives | False negatives | False positives | Recall | Precision | F1-score (SNP) |
|---|---|---|---|---|---|---|---|
| 20x | P-M-DV | 3286124 | 41371 | 475385 | 0.9876 | 0.8736 | 0.9271 |
| | Medaka | 3228058 | 59716 | 369037 | 0.9818 | 0.8974 | 0.9377 |
| | Clair | 3026716 | 300779 | 229952 | 0.9096 | 0.9294 | 0.9194 |
| 30x | P-M-DV | 3308068 | 19427 | 60871 | 0.9942 | 0.9819 | 0.9880 |
| | Medaka | 3248842 | 34884 | 59816 | 0.9894 | 0.9819 | 0.9856 |
| | Clair | 3194577 | 132918 | 121085 | 0.9601 | 0.9635 | 0.9618 |
| 40x | P-M-DV | 3312504 | 14991 | 20633 | 0.9955 | 0.9938 | 0.9947 |
| | Medaka | 3279473 | 29155 | 39141 | 0.9912 | 0.9882 | 0.9897 |
| | Clair | 3237789 | 89706 | 80265 | 0.9730 | 0.9758 | 0.9744 |
| 50x | P-M-DV | 3314808 | 12687 | 13806 | 0.9962 | 0.9959 | 0.9960 |
| | Medaka | 3298374 | 26404 | 33504 | 0.9921 | 0.9899 | 0.9910 |
| | Clair | 3254017 | 73478 | 58229 | 0.9779 | 0.9824 | 0.9802 |
| 60x | P-M-DV | 3315655 | 11840 | 12062 | 0.9964 | 0.9964 | 0.9964 |
| | Medaka | 3271294 | 24807 | 30926 | 0.9925 | 0.9906 | 0.9916 |
| | Clair | 3260364 | 67131 | 46813 | 0.9798 | 0.9858 | 0.9828 |
| 70x | P-M-DV | 3316257 | 11238 | 11217 | 0.9966 | 0.9966 | 0.9966 |
| | Medaka | 3283443 | 24010 | 28991 | 0.9927 | 0.9913 | 0.9920 |
| | Clair | 3263513 | 63982 | 39636 | 0.9808 | 0.9880 | 0.9844 |
| 80x | P-M-DV | 3316750 | 10745 | 10219 | 0.9968 | 0.9969 | 0.9969 |
| | Medaka | 3280595 | 23263 | 27321 | 0.9930 | 0.9917 | 0.9924 |
| | Clair | 3265361 | 62134 | 34616 | 0.9813 | 0.9895 | 0.9854 |
| 90x | P-M-DV | 3317032 | 10463 | 9958 | 0.9969 | 0.9970 | 0.9969 |
| | Medaka | 3293174 | 22716 | 26549 | 0.9931 | 0.9920 | 0.9926 |
| | Clair | 3266489 | 61006 | 31220 | 0.9817 | 0.9905 | 0.9861 |

Supplementary Table 3: Comparison on SNP performance between Medaka, Clair and PEPPER-Margin-DeepVariant (P-M-DV) variant callers at different coverages of HG003 sample.

| File | Read N50 | Gb | Coverage |
|---|---|---|---|
| HG001 | 21443 | 309.88 | 93.91 |
| HG002 | 50317 | 160.39 | 48.6 |
| HG003 | 44550 | 277.38 | 84.05 |
| HG004 | 47996 | 284.32 | 86.16 |
| HG005 | 49297 | 182.53 | 55.31 |
| HG006 | 50019 | 163.87 | 49.66 |
| HG007 | 50423 | 132.75 | 40.23 |

Supplementary Table 4: Sample-wise nanopore read coverage for seven Genome-In-A-Bottle (GIAB) samples.

| Sample Name | Reference | Version | Regions covered by benchmark (bp) |
|---|---|---|---|
| HG001 | GRCh37 | v3.3.2 | 2437907771 |
| HG002 | GRCh38 | v4.2.1 | 2542242843 |
| HG003 | GRCh38 | v4.2.1 | 2528531102 |
| HG004 | GRCh38 | v4.2.1 | 2524487531 |
| HG005 | GRCh37 | v3.3.2 | 2376855757 |
| HG006 | GRCh37 | v3.3.2 | 2393652163 |
| HG007 | GRCh37 | v3.3.2 | 2394471248 |

Supplementary Table 5: Details of Genome-In-A-Bottle truth set used for each genome.

| **Sample** | **Type** | **Total truth** | **True positives** | **False negatives** | **False positives** | **Recall** | **Precision** | **F1-score** |
|---|---|---|---|---|---|---|---|---|
| HG001 | SNP | 3209309 | 3203740 | 5569 | 11466 | 0.9983 | 0.9964 | 0.9973 |
| | INDEL | 481841 | 292565 | 189276 | 37718 | 0.6072 | 0.8883 | 0.7213 |
| HG003 | SNP | 3327495 | 3317032 | 10463 | 9958 | 0.9969 | 0.9970 | 0.9969 |
| | INDEL | 504501 | 303643 | 200858 | 29400 | 0.6019 | 0.9136 | 0.7257 |
| HG004 | SNP | 3346610 | 3338882 | 7728 | 7474 | 0.9977 | 0.9978 | 0.9977 |
| | INDEL | 510519 | 300258 | 210261 | 32429 | 0.5881 | 0.9046 | 0.7128 |
| HG005 | SNP | 3042623 | 3036676 | 5947 | 11807 | 0.9980 | 0.9961 | 0.9971 |
| | INDEL | 390158 | 258720 | 131438 | 30008 | 0.6631 | 0.8981 | 0.7629 |
| HG006 | SNP | 3053660 | 3047013 | 6647 | 13802 | 0.9978 | 0.9955 | 0.9967 |
| | INDEL | 394727 | 242909 | 151818 | 30518 | 0.6154 | 0.8901 | 0.7277 |
| HG007 | SNP | 3069407 | 3060423 | 8984 | 18351 | 0.9971 | 0.9940 | 0.9956 |
| | INDEL | 397103 | 236816 | 160287 | 34295 | 0.5964 | 0.8753 | 0.7094 |

Supplementary Table 6: PEPPER-Margin-DeepVariant performance on six GIAB samples with Oxford nanopore data.

| Sample name | Pipeline | Type | True positives | False negatives | False positives | Recall | Precision | F1-score |
|---|---|---|---|---|---|---|---|---|
| HG003 35x | DeepVariant | INDEL | 499277 | 5224 | 4901 | 0.9896 | 0.9907 | 0.9902 |
| | | SNP | 3323609 | 3886 | 2883 | 0.9988 | 0.9991 | 0.9990 |
| | DV-WH-DV | INDEL | 501509 | 2992 | 2935 | 0.9941 | 0.9944 | 0.9942 |
| | | SNP | 3323655 | 3840 | 2734 | 0.9988 | 0.9992 | 0.9990 |
| | DV-M-DV | INDEL | 501567 | 2934 | 2746 | **0.9942** | **0.9948** | **0.9945** |
| | | SNP | 3323586 | 3909 | 1841 | **0.9988** | **0.9994** | **0.9991** |
| | P-M-DV | INDEL | 501539 | 2962 | 2816 | 0.9941 | 0.9946 | 0.9944 |
| | | SNP | 3323607 | 3888 | 2501 | 0.9988 | 0.9992 | 0.9990 |
| HG004 35x | DeepVariant | INDEL | 504939 | 5580 | 5217 | 0.9891 | 0.9902 | 0.9896 |
| | | SNP | 3343142 | 3468 | 2274 | 0.9990 | 0.9993 | 0.9991 |
| | DV-WH-DV | INDEL | 507288 | 3231 | 2966 | 0.9937 | 0.9944 | 0.9940 |
| | | SNP | 3343074 | 3536 | 1771 | 0.9989 | 0.9995 | 0.9992 |
| | DV-M-DV | INDEL | 507351 | 3168 | 2846 | **0.9938** | **0.9946** | **0.9942** |
| | | SNP | 3342966 | 3644 | 1491 | **0.9989** | **0.9996** | **0.9992** |
| | P-M-DV | INDEL | 507313 | 3206 | 2903 | 0.9937 | 0.9945 | 0.9941 |
| | | SNP | 3342928 | 3682 | 1721 | 0.9989 | 0.9995 | 0.9992 |

Supplementary Table 8: PacBio HiFi variant calling perfomance comparison between PEPPER-Margin-DeepVariant (P-M-DV), DeepVariant-WhatsHap-DeepVariant (DV-WH-DV), DeepVariant-Margin-DV (DV-M-DV), DeepVariant only.

| Sample | Pipeline | SNP calling runtime | Phasing runtime | Variant calling runtime | Total runtime | Cost |
|---|---|---|---|---|---|---|
| HG003 35x PacBio HiFi | DeepVariant Whatshap DeepVariant | 03:48:45 (n1-std-96) | 07:32:35 (n1-std-2) | 03:50:32 (n1-std-96) | 15:11:52 | $36.24 |
| | DeepVariant Margin DeepVariant | 03:48:45 (n1-std-96) | 00:24:51 (n1-std-96) | 03:50:07 (n1-std-96) | 08:03:43 | $36.71 |
| | PEPPER Margin DeepVariant | 1:28:49 (n1-std-96) | 00:23:25 (n1-std-96) | 4:03:14 (n1-std-96) | 05:55:28 | $26.99 |
| HG004 35x PacBio HiFi | DeepVariant Whatshap DeepVariant | 04:03:58 (n1-std-96) | 07:44:56 (n1-std-2) | 03:58:35 (n1-std-96) | 15:47:29 | $37.35 |
| | DeepVariant Margin DeepVariant | 04:03:58 (n1-std-96) | 00:26:09 (n1-std-96) | 03:56:29 (n1-std-96) | 08:26:36 | $38.45 |
| | PEPPER Margin DeepVariant | 1:25:03 (n1-std-96) | 0:23:41 (n1-std-96) | 4:10:13 (n1-std-96) | 05:58:57 | $27.26 |

Supplementary Table 9: PacBio HiFi variant calling run-time comparison between three haplotype-aware pipelines on 35× coverage HG003 and HG004 samples. We used $4.56/h `n1-standard-96 (n1-std-96)` and $0.09/h `n1-standard-2 (n1-std-2)` instance types on google cloud platform for this analysis.

| Sample | Variant type | Sequencing technology | True positives | False negatives | False positives | Recall | Precision | F1-score |
|---|---|---|---|---|---|---|---|---|
| HG003 | SNP | Nanopore | 3317032 | 10463 | 9958 | 0.9969 | 0.9970 | 0.9969 |
| | | Illumina | 3307988 | 19508 | 4808 | 0.9941 | 0.9985 | 0.9963 |
| | | PacBio HiFi | 3323607 | 3888 | 2501 | 0.9988 | 0.9992 | 0.9990 |
| | INDEL | Nanopore | 303643 | 200858 | 29400 | 0.6019 | 0.9136 | 0.7257 |
| | | Illumina | 501546 | 2955 | 1276 | 0.9941 | 0.9976 | 0.9959 |
| | | PacBio HiFi | 501539 | 2962 | 2816 | 0.9941 | 0.9946 | 0.9944 |
| HG004 | SNP | Nanopore | 3338882 | 7728 | 7474 | 0.9977 | 0.9978 | 0.9977 |
| | | Illumina | 3326040 | 20570 | 4476 | 0.9939 | 0.9987 | 0.9962 |
| | | PacBio HiFi | 3342928 | 3682 | 1721 | 0.9989 | 0.9995 | 0.9992 |
| | INDEL | Nanopore | 300258 | 210261 | 32429 | 0.5881 | 0.9046 | 0.7128 |
| | | Illumina | 507418 | 3101 | 1284 | 0.9939 | 0.9976 | 0.9958 |
| | | PacBio HiFi | 507313 | 3206 | 2903 | 0.9937 | 0.9945 | 0.9941 |

Supplementary Table 10: Variant calling performance comparison in all benchmark regions between Oxford Nanopore Technology (ONT), Illumina NovaSeq (Illumina) and PacBio HiFi sequencing technology. Illumina variant calls are generated with DeepVariant v1.1 and ONT and PacBio HiFi variant calls are generated with PEPPER-Margin-DeepVariant.

| Region | Sample | Platform | Total truth | True positives | False negatives | False positives | Recall | Precision | F1-score |
|---|---|---|---|---|---|---|---|---|---|
| MHC (SNP) | HG003 | ONT | 19543 | 19437 | 106 | 56 | 0.9946 | 0.9971 | 0.9958 |
| | | Illumina | 19544 | 19336 | 208 | 31 | 0.9894 | 0.9984 | 0.9939 |
| | | PacBio | 19543 | 19391 | 152 | 39 | 0.9922 | 0.9980 | 0.9951 |
| | HG004 | ONT | 19271 | 19181 | 90 | 42 | 0.9953 | 0.9978 | 0.9966 |
| | | Illumina | 19271 | 18998 | 273 | 31 | 0.9858 | 0.9984 | 0.9921 |
| | | PacBio | 19271 | 19112 | 159 | 12 | 0.9917 | 0.9994 | 0.9955 |
| Seg. Dup. (SNP) | HG003 | ONT | 121960 | 119838 | 2122 | 2288 | 0.9826 | 0.9813 | 0.9819 |
| | | Illumina | 121960 | 112293 | 9667 | 2905 | 0.9207 | 0.9748 | 0.9470 |
| | | PacBio | 121960 | 119003 | 2957 | 818 | 0.9758 | 0.9932 | 0.9844 |
| | HG004 | ONT | 122191 | 120107 | 2084 | 1693 | 0.9829 | 0.9861 | 0.9845 |
| | | Illumina | 122191 | 112296 | 9895 | 2710 | 0.9190 | 0.9764 | 0.9469 |
| | | PacBio | 122191 | 119713 | 2478 | 672 | 0.9797 | 0.9944 | 0.9870 |
| Low map. (SNP) | HG003 | ONT | 192520 | 190380 | 2140 | 2152 | 0.9889 | 0.9888 | 0.9889 |
| | | Illumina | 192520 | 174763 | 17757 | 3627 | 0.9078 | 0.9797 | 0.9423 |
| | | PacBio | 192520 | 189453 | 3067 | 888 | 0.9841 | 0.9953 | 0.9897 |
| | HG004 | ONT | 192653 | 190671 | 1982 | 1634 | 0.9897 | 0.9915 | 0.9906 |
| | | Illumina | 192653 | 174196 | 18457 | 3510 | 0.9042 | 0.9803 | 0.9407 |
| | | PacBio | 192653 | 190118 | 2535 | 653 | 0.9868 | 0.9966 | 0.9917 |
| 250bp+ non-unique (SNP) | HG003 | ONT | 13608 | 12594 | 1014 | 592 | 0.9255 | 0.9552 | 0.9401 |
| | | Illumina | 13608 | 7420 | 6188 | 1377 | 0.5453 | 0.8436 | 0.6624 |
| | | PacBio | 13608 | 11613 | 1995 | 380 | 0.8534 | 0.9684 | 0.9072 |
| | HG004 | ONT | 13492 | 12615 | 877 | 413 | 0.9350 | 0.9683 | 0.9514 |
| | | Illumina | 13492 | 7235 | 6257 | 1323 | 0.5362 | 0.8455 | 0.6563 |
| | | PacBio | 13492 | 11847 | 1645 | 284 | 0.8781 | 0.9766 | 0.9247 |

Supplementary Table 11: SNP performance in difficult-to-map regions with Illumina, PacBio HiFi and Oxford nanopore data.

| Region | Sample | Platform | Total truth | True positives | False negatives | False positives | Recall | Precision | F1-score |
|---|---|---|---|---|---|---|---|---|---|
| H.poly. (7bp-11bp) | HG003 | ONT | 70736 | 68620 | 2116 | 2464 | 0.9701 | 0.9654 | 0.9677 |
| | | Illumina | 70737 | 70632 | 105 | 60 | 0.9985 | 0.9992 | 0.9988 |
| | | PacBio | 70736 | 70645 | 91 | 93 | 0.9987 | 0.9987 | 0.9987 |
| | HG004 | ONT | 71141 | 68912 | 2229 | 2665 | 0.9687 | 0.9628 | 0.9657 |
| | | Illumina | 71141 | 71045 | 96 | 39 | 0.9987 | 0.9995 | 0.9991 |
| | | PacBio | 71142 | 71032 | 110 | 127 | 0.9985 | 0.9982 | 0.9983 |
| H.Poly. 11bp+ | HG003 | ONT | 12187 | 10708 | 1479 | 1359 | 0.8786 | 0.8879 | 0.8832 |
| | | Illumina | 12188 | 12176 | 12 | 16 | 0.9990 | 0.9987 | 0.9989 |
| | | PacBio | 12187 | 12005 | 182 | 203 | 0.9851 | 0.9841 | 0.9846 |
| | HG004 | ONT | 12494 | 10751 | 1743 | 1441 | 0.8605 | 0.8823 | 0.8713 |
| | | Illumina | 12494 | 12478 | 16 | 38 | 0.9987 | 0.9971 | 0.9979 |
| | | PacBio | 12494 | 12332 | 162 | 182 | 0.9870 | 0.9861 | 0.9866 |
| Di-Mer repeat (11bp-50bp) | HG003 | ONT | 18817 | 18322 | 495 | 668 | 0.9737 | 0.9654 | 0.9695 |
| | | Illumina | 18817 | 18778 | 39 | 34 | 0.9979 | 0.9982 | 0.9981 |
| | | PacBio | 18817 | 18763 | 54 | 120 | 0.9971 | 0.9939 | 0.9955 |
| | HG004 | ONT | 18925 | 18417 | 508 | 676 | 0.9732 | 0.9652 | 0.9692 |
| | | Illumina | 18925 | 18880 | 45 | 41 | 0.9976 | 0.9979 | 0.9978 |
| | | PacBio | 18925 | 18873 | 52 | 109 | 0.9973 | 0.9945 | 0.9959 |
| Tri-Mer repeat (15bp-50bp) | HG003 | ONT | 4179 | 4129 | 50 | 89 | 0.9880 | 0.9791 | 0.9835 |
| | | Illumina | 4179 | 4172 | 7 | 3 | 0.9983 | 0.9993 | 0.9988 |
| | | PacBio | 4179 | 4153 | 26 | 22 | 0.9938 | 0.9948 | 0.9943 |
| | HG004 | ONT | 4213 | 4175 | 38 | 92 | 0.9910 | 0.9785 | 0.9847 |
| | | Illumina | 4213 | 4210 | 3 | 2 | 0.9993 | 0.9995 | 0.9994 |
| | | PacBio | 4213 | 4196 | 17 | 18 | 0.9960 | 0.9958 | 0.9959 |

Supplementary Table 12: SNP performance in low-complexity regions with Illumina, PacBio HiFi and Oxford nanopore data.

| Data | Tool | Phased Variants | Unphased Variants | Blocks | Median Variants per Block | Average Variants per Block | Median BP per Block | Average BP per Block | Block N50 |
|---|---|---|---|---|---|---|---|---|---|
| ONT 25x | Margin | 2293009 | 1008276 | 2536 | 347 | 904 | 503808 | 1056597 | 2067806 |
| | WhatsHap | 2452395 | 849215 | 2297 | 395 | 1068 | 523694 | 1177941 | 2372651 |
| ONT 50x | Margin | 2275697 | 875317 | 1376 | 613 | 1654 | 853602 | 1993709 | 4211518 |
| | WhatsHap | 2391670 | 759715 | 1172 | 822 | 2041 | 1049089 | 2355537 | 4900234 |
| ONT 75x | Margin | 2091713 | 1023259 | 1167 | 496 | 1792 | 769148 | 2372510 | 6126250 |
| | WhatsHap | 2393421 | 722297 | 812 | 955 | 2948 | 1167915 | 3430964 | 8266083 |
| HiFi 35x | Margin | 2327420 | 1035855 | 14069 | 15 | 165 | 48362 | 154095 | 242226 |
| | WhatsHap | 2412900 | 954503 | 14061 | 14 | 172 | 48362 | 155745 | 252972 |

Supplementary Table 13: Details of Margin and WhatsHap phasing output on HG001 sample with Oxford Nanopore (ONT) and PacBio HiFi data. Results are generated with `whatshap stats` command.

| Data | Tool | Data | Tool | Assessed Pairs | Switches | Switch Rate | Hamming | Hamming Rate |
|---|---|---|---|---|---|---|---|---|
| ONT 25x | Margin | ONT 25x | Margin | 1901418 | 16639 | 0.00875 | 162341 | 0.0854 |
| | WhatsHap | ONT 25x | WhatsHap | 1917571 | 17696 | 0.00923 | 177660 | 0.0926 |
| ONT 50x | Margin | ONT 50x | Margin | 1895721 | 16252 | 0.00857 | 195897 | 0.1033 |
| | WhatsHap | ONT 50x | WhatsHap | 1926257 | 17513 | 0.00909 | 161079 | 0.0836 |
| ONT 75x | Margin | ONT 75x | Margin | 1759253 | 14356 | 0.00816 | 174052 | 0.0989 |
| | WhatsHap | ONT 75x | WhatsHap | 1927665 | 17462 | 0.00906 | 179655 | 0.0932 |
| HiFi 35x | Margin | HiFi 35x | Margin | 1908770 | 17077 | 0.00895 | 24187 | 0.0127 |
| | WhatsHap | HiFi 35x | WhatsHap | 1914368 | 17801 | 0.00930 | 28973 | 0.0151 |

Supplementary Table 14: Comparison of Margin and WhatsHap phasesets of HG001 sample with Oxford Nanopore (ONT) and PacBio HiFi data. Comparison is performed with `whatshap compare` command.

| Data | Tool | Average Accuracy | Average Reads per 10kb | Average Tagged Reads per 10kb |
|---|---|---|---|---|
| ONT 55x Chr1 | Margin | 95.57 | 63x | 58x |
| | WhatsHap | 95.34 | 63x | 58x |
| CCS 35x Chr1 | Margin | 97.64 | 39x | 36x |
| | WhatsHap | | | |

Supplementary Table 15: Haplotagging Results

| Data | Tool | Module | Max Threads | Max Memory | Runtime (min) | Instance Type | Instance Cost ($/hr) | Cost ($) |
|---|---|---|---|---|---|---|---|---|
| ONT 25x | Margin | Haplotag | 64 | 20 | 21 | n1-highcpu-64 | 2.267 | 0.79 |
| | | Phase VCF | 64 | 15 | 15 | n1-highcpu-64 | 2.267 | 0.56 |
| | | Total | – | – | 36 | n1-highcpu-64 | 2.267 | 1.36 |
| | WhatsHap | Phase | 2 | 3 | 347 | n1-standard-2 | 0.095 | 0.54 |
| | | Haplotag | 2 | 3 | 247 | n1-standard-2 | 0.095 | 0.39 |
| | | Total | – | – | 941 | n1-standard-2 | 0.095 | 1.48 |
| ONT 50x | Margin | Haplotag | 64 | 28 | 54 | n1-highcpu-64 | 2.267 | 2.04 |
| | | Phase VCF | 64 | 18 | 30 | n1-highcpu-64 | 2.267 | 1.13 |
| | | Total | – | – | 84 | n1-highcpu-64 | 2.267 | 3.17 |
| | WhatsHap | Phase | 2 | 3 | 446 | n1-standard-2 | 0.095 | 0.7 |
| | | Haplotag | 2 | 3 | 444 | n1-standard-2 | 0.095 | 0.7 |
| | | Total | – | – | 1336 | n1-standard-2 | 0.095 | 2.11 |
| ONT 75x | Margin | Haplotag | 64 | 35 | 80 | n1-highcpu-64 | 2.267 | 3.02 |
| | | Phase VCF | 64 | 22 | 43 | n1-highcpu-64 | 2.267 | 1.62 |
| | | Total | – | – | 123 | n1-highcpu-64 | 2.267 | 4.64 |
| | WhatsHap | Phase | 2 | 3 | 522 | n1-standard-2 | 0.095 | 0.82 |
| | | Haplotag | 2 | 3 | 644 | n1-standard-2 | 0.095 | 1.01 |
| | | Total | – | – | 1688 | n1-standard-2 | 0.095 | 2.67 |
| HiFi 35x | Margin | Haplotag | 64 | 19 | 19 | n1-highcpu-64 | 2.267 | 0.71 |
| | | Phase VCF | 64 | 18 | 14 | n1-highcpu-64 | 2.267 | 0.52 |
| | | Total | – | – | 33 | n1-highcpu-64 | 2.267 | 1.24 |
| | WhatsHap | Phase | 2 | 3 | 277 | n1-standard-2 | 0.095 | 0.43 |
| | | Haplotag | 2 | 3 | 210 | n1-standard-2 | 0.095 | 0.33 |
| | | Total | – | – | 764 | n1-standard-2 | 0.095 | 1.2 |

Supplementary Table 16: Margin/WhatsHap Runtimes. Total runtimes are sum of Haplotag and Phase VCF runtimes for Margin, and sum of 2x Phase and 1x Haplotag for WhatsHap, as `whatshap haplotag` requires a phased VCF.

| Sample | Assembler | Polisher | Assembly haplotype | NG50 | Estimated QV YAK (k=31) | Switch error rate | Hamming error |
|---|---|---|---|---|---|---|---|
| HG005 | Flye | - | Haploid | 37254637 | 31.08 | 0.146333 | 0.319502 |
| | Shasta | - | Haploid | 39831103 | 32 | 0.16431 | 0.293283 |
| | | P-M-DV (ONT) | HP-1 | 39820763 | 35.06 | 0.058178 | 0.207221 |
| | | | HP-2 | 39820481 | 35.06 | 0.059199 | 0.218271 |
| | | P-M-DV (PacBio HiFi) | HP-1 | 39808277 | 43.54 | 0.028165 | 0.26687 |
| | | | HP-2 | 39809097 | 43.5 | 0.028253 | 0.264903 |
| | Trio-hifiasm | - | mat | 51324672 | 51.81 | 0.007056 | 0.009601 |
| | | | pat | 50669010 | 51.72 | 0.003106 | 0.004542 |
| HG00733 | Flye | - | Haploid | 36602095 | 31.93 | 0.226708 | 0.455478 |
| | Shasta | - | Haploid | 42512208 | 32.7 | 0.263731 | 0.4387 |
| | | P-M-DV (ONT) | HP-1 | 42497702 | 35.83 | 0.09903 | 0.319373 |
| | | | HP-2 | 42498275 | 35.84 | 0.098502 | 0.320807 |
| | | P-M-DV (PacBio HiFi) | HP-1 | 42475072 | 43.83 | 0.050551 | 0.401146 |
| | | | HP-2 | 42476106 | 43.85 | 0.049385 | 0.406129 |
| | Trio-hifiasm | - | mat | 32479553 | 53.6 | 0.0102 | 0.012044 |
| | | | pat | 35318917 | 53.35 | 0.010144 | 0.010069 |
| HG02723 | Flye | - | Haploid | 39652856 | 31.88 | 0.24692 | 0.454764 |
| | Shasta | - | Haploid | 49185987 | 32.52 | 0.28754 | 0.424615 |
| | | P-M-DV (ONT) | HP-1 | 49165039 | 35.8 | 0.104264 | 0.248018 |
| | | | HP-2 | 49164831 | 35.79 | 0.103455 | 0.238674 |
| | | P-M-DV (PacBio HiFi) | HP-1 | 49146102 | 43.46 | 0.046367 | 0.365246 |
| | | | HP-2 | 49143792 | 43.38 | 0.046215 | 0.363784 |
| | Trio-hifiasm | - | mat | 19737990 | 56.27 | 0.005794 | 0.007677 |
| | | | pat | 22214675 | 55.94 | 0.006683 | 0.009111 |

Supplementary Table 17: Diploid assembly polishing results of PEPPER-Margin-DeepVariant (P-M-DV) pipeline on HG005, HG00733 and HG02723 samples. We report estimated quality value (QV), switch error rate and hamming error using YAK assembly assessment tool.

| Sample | Assembler | Polisher | Estimated QV YAK (k=31) |
|---|---|---|---|
| CHM13 chrX | Flye | - | 32.85 |
| | Shasta | - | 34.601 |
| | | P-M-DV (ONT) | 36.91 |
| | | P-M-DV (PacBio HiFi) | 42.765 |
| | Hifiasm | - | 53.039 |

Supplementary Table 18: Haploid assembly polishing results of PEPPER-Margin-DeepVariant (P-M-DV) pipeline on CHM13-chrX. We report estimated quality value (QV) using YAK assembly assessment tool.

| Type | Data | Gene Type | Subset | Recall | Precision | F1 Score |
|---|---|---|---|---|---|---|
| SNP | Nanopore | all regions | all regions | 0.998169 | 0.996314 | 0.997241 |
| | | all genes | all genes | 0.998097 | 0.996481 | 0.997289 |
| | | protein coding | all cds | 0.998641 | 0.997887 | 0.998263 |
| | | | all exons | 0.998158 | 0.997675 | 0.997916 |
| | | | all genes | 0.99799 | 0.996751 | 0.99737 |
| | PacBio HiFi | all regions | all regions | 0.999391 | 0.998062 | 0.998726 |
| | | all genes | all genes | 0.999384 | 0.998197 | 0.99879 |
| | | protein coding | all cds | 0.999446 | 0.998994 | 0.99922 |
| | | | all exons | 0.999502 | 0.999117 | 0.99931 |
| | | | all genes | 0.999382 | 0.998441 | 0.998912 |
| INDEL | Nanopore | all regions | all regions | 0.60077 | 0.878512 | 0.713567 |
| | | all genes | all genes | 0.595042 | 0.877943 | 0.709325 |
| | | protein coding | all cds | 0.799544 | 0.926893 | 0.858522 |
| | | | all exons | 0.632435 | 0.896594 | 0.741696 |
| | | | all genes | 0.584914 | 0.876731 | 0.701692 |
| | PacBio HiFi | all regions | all regions | 0.948736 | 0.92602 | 0.937241 |
| | | all genes | all genes | 0.947847 | 0.922887 | 0.935201 |
| | | protein coding | all cds | 0.984055 | 0.909278 | 0.94519 |
| | | | all exons | 0.955149 | 0.927624 | 0.941186 |
| | | | all genes | 0.946128 | 0.918991 | 0.932362 |

Supplementary Table 19: Accuracy stats for ONT and CCS calls made on GRCh37 with HG001 data in high confidence regions against GIAB v3.3.2 stratified by all gene and protein coding gene, further stratified by whole gene, exon, CDS as annotated by GENCODE v35lift37. CDS regions are coding sequences, and include start and stop codons for this analysis.

| Gene Region | Subset | Subset Size | Subset High Confidence Size | High Confidence Ratio | High Confidence Whole Genome Ratio |
|---|---|---|---|---|---|
| Genome | – | 2951332653 | 2579466415 | 0.874001 | 0.874001 |
| All Genes | – | 1982798080 | 1591767788 | 0.802789 | 0.539339 |
| Protein Coding | Coding Sequence | 114906140 | 31986772 | 0.278373 | 0.010838 |
| Protein Coding | Exon | 283314507 | 92457254 | 0.326341 | 0.031327 |
| Protein Coding | Gene Regions | 1367165648 | 1201166019 | 0.878581 | 0.406991 |

Supplementary Table 20: Size of GENCODE Gene Regions

| Phasing Coverage | Switch Errors Present | SNP, INDEL Errors | Gene Region | Count | 25 Quartile Gene Size | Median Gene Size | 75 Quartile Gene Size |
|---|---|---|---|---|---|---|---|
| wholly | no error | no error | gene | 1738 | 1438 | 3332 | 8005 |
| | | | exon | 3121 | 2845 | 10303 | 34848 |
| | | | cds | 3481 | 3163 | 11478 | 39123 |
| | | error | gene | 1764 | 15676 | 35729 | 80335 |
| | | | exon | 381 | 7533 | 27218 | 67507 |
| | | | cds | 21 | 3161 | 5190 | 31102 |
| | error | no error | gene | 15 | 1685 | 4868 | 15383 |
| | | | exon | 33 | 8703 | 19682 | 86981 |
| | | | cds | 37 | 8703 | 20549 | 72326 |
| | | error | gene | 23 | 18072 | 63546 | 107182 |
| | | | exon | 5 | 7680 | 47627 | 63546 |
| | | | cds | 1 | 7680 | 7680 | 7680 |
| partially | no error | no error | gene | 6 | 4760 | 10482 | 36920 |
| | | | exon | 29 | 25339 | 102612 | 147661 |
| | | | cds | 37 | 25339 | 102612 | 161893 |
| | | error | gene | 31 | 44745 | 110698 | 223634 |
| | | | exon | 8 | 54807 | 99792 | 318869 |
| | | | cds | 0 | – | – | – |
| | error | all | all | 0 | – | – | – |
| not | – | no error | gene | 125 | 1769 | 3060 | 8819 |
| | | | exon | 201 | 2352 | 8744 | 29504 |
| | | | cds | 214 | 2478 | 9365 | 29958 |
| | | error | gene | 91 | 19316 | 33429 | 69811 |
| | | | exon | 15 | 10690 | 23836 | 35712 |
| | | | cds | 2 | 44794 | 53073 | 61351 |

Supplementary Table 21: Gencode protein coding genes with coding sequence (CDS, start_codon, and stop_codon) 80% spanned by high confidence stratified by how phased it is by Margin, whether there were switch errors, whether there were SNP or INDEL errors, and gene region for HG001 with 75x Nanopore data on GRCh37. Three gene length quartiles are presented for the groupings.

| Phasing Coverage | Switch Errors Present | SNP, INDEL Errors | Gene Region | Count | 25 Quartile Gene Size | Median Gene Size | 75 Quartile Gene Size |
|---|---|---|---|---|---|---|---|
| wholly | no error | no error | gene | 2086 | 2184 | 5907 | 17294 |
| | | | exon | 2446 | 2586 | 8471 | 27219 |
| | | | cds | 2474 | 2615 | 8536 | 27351 |
| | | error | gene | 390 | 23230 | 51309 | 102861 |
| | | | exon | 30 | 7098 | 15068 | 65957 |
| | | | cds | 2 | 60929 | 108132 | 155335 |
| | error | no error | gene | 18 | 1690 | 6798 | 11879 |
| | | | exon | 23 | 2321 | 8703 | 23001 |
| | | | cds | 24 | 2567 | 9247 | 19731 |
| | | error | gene | 6 | 13297 | 23861 | 43535 |
| | | | exon | 1 | 12242 | 12242 | 12242 |
| | | | cds | 0 | – | – | – |
| partially | no error | no error | gene | 190 | 13338 | 34319 | 67099 |
| | | | exon | 354 | 28087 | 68206 | 146543 |
| | | | cds | 360 | 28289 | 68510 | 145331 |
| | | error | gene | 170 | 76695 | 137702 | 255101 |
| | | | exon | 6 | 55573 | 75267 | 104898 |
| | | | cds | 0 | – | – | – |
| | error | no error | gene | 2 | 20915 | 21281 | 21647 |
| | | | exon | 5 | 22014 | 109387 | 127176 |
| | | | cds | 5 | 22014 | 109387 | 127176 |
| | | error | gene | 3 | 118281 | 127176 | 138247 |
| | | | exon | 0 | – | – | – |
| | | | cds | 0 | – | – | – |
| not | no error | no error | gene | 741 | 2565 | 8419 | 23809 |
| | | | exon | 917 | 3533 | 13187 | 43784 |
| | | | cds | 928 | 3639 | 13351 | 43785 |
| | | error | gene | 187 | 30101 | 75166 | 156763 |
| | | | exon | 11 | 10504 | 29953 | 68614 |
| | | | cds | 0 | – | – | – |

Supplementary Table 22: Gencode protein coding genes with coding sequence (CDS, start_codon, and stop_codon) 80% spanned by high confidence stratified by how phased it is by Margin, whether there were switch errors, whether there were SNP or INDEL errors, and gene region for HG001 with 35x PacBio HiFi data on GRCh37. Three gene length quartiles are presented for the groupings.
